# Mechanisms of Ozone Effects on Plant Stress in Soybean Across Growing Season: From Leaf to Regional Perspective

**DOI:** 10.1101/2025.09.29.679296

**Authors:** Luka Mamić, Mj Riches, Rose K. Rossell, Delphine K. Farmer

## Abstract

Ground-level ozone (O₃) is a major constraint on agricultural productivity, yet most knowledge comes from controlled fumigation experiments using chronic exposures that differ from the episodic conditions crops experience in the field. Here, we combine a five-week chamber experiment with multi-year satellite observations (2018-2021, Arkansas, U.S.) to investigate how O₃ affects photosynthesis, efficiency, and growth across scales of soybean plants (*Glycine max*). At the leaf level, initial O₃ fumigation (80 ppb for 4 h) caused the strongest suppression of CO₂ assimilation (A), stomatal conductance (Gs), and photosystem II efficiency (ΦPSII), indicating entry into a physiological strain phase. Recovery between exposures was incomplete, leading to sustained growth reductions despite moderate O₃ levels. At the regional scale, analysis of solar-induced fluorescence (SIF) and MODIS productivity metrics revealed parallel patterns. Early-season O₃ episodes produced greater suppression of SIF, GPP, and Gs compared to equivalent late-season events, and recovery lagged for several weeks. Seasonal yield proxies were best explained not by total O₃ accumulation, but by early- and peak-season exposures, which accounted for up to 98% of variance across four growing seasons. Our findings highlight that the timing of O₃ episodes is more consequential than cumulative dose, and that functional indicators such as SIF can detect strain-phase stress before structural indices diverge. By linking controlled experiments with regional-scale satellite monitoring, this study advances mechanistic understanding of O₃ impacts on soybean and supports the development of remote sensing–based early warning tools for crop management.

## 1. Introduction

Air pollution threatens agricultural productivity and food security around the globe (Dong & Wang, 2023; Tai et al., 2014). Among air pollutants, ground-level ozone (O₃) is one of the most important stressors to global crop production (Ashmore, 2005; Emberson et al., 2009). O₃ is a secondary air pollutant formed via photochemical reactions between nitrogen oxides and volatile organic compounds. The terrestrial biosphere is a key sink for O₃, which may reduce atmospheric concentrations of this pollutant, but means that O₃ deposition can directly impact crop yield and productivity (Clifton et al., 2020; Fares et al., 2014). Even short-term periods of elevated O₃ can significantly decrease yields (Mills, Pleijel, et al., 2018; Tai et al., 2014), with soybean being one of the most sensitive – yet economically important – crops (Betzelberger et al., 2012; Morgan et al., 2003).

While there is a broad understanding of O₃-plant interactions, many details remain unclear, such as species-specific physiological responses (Feng et al., 2015), the mechanisms underlying photosynthetic and metabolic impairments (Ainsworth, 2017), and interactions with climatic and agronomic factors (Mills, Sharps, et al., 2018). Plants primarily regulate O₃ entry through stomatal control, yet stomatal uptake accounts for only ∼45% of total ecosystem-level O₃ deposition, with the remainder occurring via non-stomatal pathways such as uptake by cuticular surfaces, soil, and wet leaf films (Clifton et al., 2020). These non-stomatal routes are especially important in dry conditions when stomata are partially closed, as harmful oxidative stress may persist internally (Wong et al., 2022), eventually impairing photosynthesis and accelerating senescence which leads to yield losses in sensitive crops (McGrath et al., 2015; Morgan et al., 2003). Felzer et al. (2007) used a coupled biosphere-atmosphere model to estimate that, under extreme conditions, surface O₃ could reduce global crop yields by up to 20% for wheat and 60% for soybeans by 2100. These reductions stem from the chronic suppression of photosynthesis and biomass accumulation based on dose-response functions from chamber and field experiments built into their model. When these losses were integrated over time and scaled to global agricultural output, the cumulative economic loss was estimated to be up to $8 trillion. This loss would primarily affect major food-producing regions in North America and Asia. Despite recognition of this scale, most large-scale estimates still rely on simplified exposure-response functions (Feng & Kobayashi, 2009; Leung et al., 2022; Mills, Sharps, et al., 2018; Montes et al., 2022; Tai et al., 2021), while the mechanistic pathways of O₃ damage across growth stages and environments remain poorly resolved. In particular, little is known about how repeated, realistic O₃ exposures interact with climatic variability to shape the transition from hidden physiological strain to visible damage and yield loss.

Plant stress can be generally described by three successive phases: (i) an initial state of applied force, (ii) a strain phase in which stress is expressed before visible damage occurs, and (iii) the damage phase, in which acute and chronic injury become visible on the leaf surface (Meroni et al., 2009). These phases are particularly challenging to study under O₃ exposure because they require carefully controlled fumigation with a highly reactive and toxic gas. Many experiments have relied on either a single acute pulse of unrealistically high O₃ (Chen et al., 2009; Singh et al., 2009) or long-term chronic exposures (Morgan et al., 2003; Osborne et al., 2016). In contrast, O₃ in the field fluctuates with the diurnal cycle and is punctuated by episodic high-O₃ days (Clifton et al., 2020; Kavassalis & Murphy, 2017). The physiological effects of repeated exposures to realistic, fluctuating concentrations may therefore differ substantially from those observed under strictly acute or chronic treatments.

Soybean, the second most widely grown crop in the United States (U.S.) and a supplier of more than one-third of global production (Liu & Desai, 2021), is highly susceptible to elevated O₃, which significantly reduces photosynthesis, stomatal conductance (Gs), biomass accumulation, and seed yield (Betzelberger et al., 2012; Morgan et al., 2003; Singh et al., 2009). Meta-analyses report average shoot biomass reductions of up to 34% and yield reductions of ∼24% under chronic exposures near 70 parts per billion by volume (ppb) O₃ (Morgan et al., 2003). The main physiological processes affected by O_3_ include carboxylation efficiency (the capacity to fix CO₂ during photosynthesis), photosystem II efficiency (ΦPSII, the effectiveness of light-driven electron transport), and carbon allocation patterns (the distribution of assimilated carbon between growth and storage) (Betzelberger et al., 2012; Chen et al., 2009). Variations among cultivars and climatic conditions further complicate these responses (Osborne et al., 2016).

At the field scale, the SoyFACE (Soybean Free Air Concentration Enrichment) project provided critical evidence on the effects of chronic O₃ under open-air, agronomic conditions. Experiments at SoyFACE demonstrated significant reductions in canopy photosynthesis, earlier leaf senescence, and lower harvest index values in soybeans exposed to elevated O₃ (∼60-70 ppb average daytime concentration, sustained across full growing seasons) (Ainsworth et al., 2014; Eastburn et al., 2010; Feng & Kobayashi, 2009; Montes et al., 2022; Wu et al., 2024). These physiological effects were often undetectable with canopy greenness-based indices such as the normalized difference vegetation index (NDVI), which primarily captures structural leaf area and chlorophyll content. By contrast, solar-induced chlorophyll fluorescence (SIF), a direct proxy for photochemical efficiency, and gas exchange measurements of photosynthesis and Gs provide more sensitive indicators of early stress. Using these approaches, SoyFACE studies identified the strain phase of O₃ stress (Wu et al., 2024). Furthermore, SoyFACE results confirmed reproductive growth stages are especially vulnerable, showing reduced maximum rate of carboxylation (Vcmax), maximum rate of electron transport (Jmax), and faster canopy decline under elevated O₃ conditions (Ainsworth et al., 2014). While SoyFACE and similar experiments provide key insight into soybean responses, they simulated future chronic air quality scenarios by maintaining elevated O₃ for entire growing seasons. In contrast, agricultural ecosystems today are often exposed to O₃ in shorter, episodic bursts, such as a few hours on high-O₃ afternoons, or multiple exceedance events across a season depending on location (Chang et al., 2025; Cooper et al., 2024; Liu & Desai, 2021). Defining what constitutes “high O₃” in agronomic terms, and understanding how repeated realistic exposures differ from strictly acute or strictly chronic treatments, remains an unresolved challenge. These differences are critical for establishing thresholds for O₃ damage in soybean (Feng & Kobayashi, 2009; Mills, Sharps, et al., 2018) and for clarifying the mechanisms by which plants defend against and recover from varying O₃ stress regimes (Chen et al., 2009; Osborne et al., 2016).

Remote sensing offers a useful pathway to investigate these knowledge gaps across agroecosystems. However, traditional remote sensing indices (e.g., NDVI) can only detect the damage phase of stress after visible morphological changes have occurred (Mahlein et al., 2012; Pinter et al., 2003). At that stage, damage is often irreversible, as photosynthetic machinery and leaf area are already degraded and yield potential cannot be recovered (Ainsworth et al., 2014; Morgan et al., 2003). However, if the stressor is removed during the strain phase, the plant can partially recover and reestablish a new physiological standard (Meroni et al., 2009). SIF is a promising parameter for detecting the strain phase earlier. Ground-based SIF measurements can monitor photosynthetic efficiency before visible symptoms develop (Wu et al., 2024). Recently-available satellite SIF data from, for example, the TROPOspheric Monitoring Instrument (TROPOMI) onboard Sentinel-5P, provides an opportunity to identify crop stress across larger scales and in near-real time. Such analyses, however, require a mechanistic understanding of how crop fluorescence responds to O_3_ exposure.

This study addresses how repeated, realistic O₃ exposures interact with environmental conditions to affect soybean physiology, and the need to evaluate when and how such stress becomes detectable with remote sensing. We combine a five-week plant-level exposure experiment with multi-year satellite observations of soybean fields in Arkansas, U.S. (2018-2021) to investigate O₃ stress across spatial and temporal scales. Specifically, we aim to (i) identify the strain phase of O₃ stress, when physiological impairment occurs before visible damage; (ii) determine when during the growing season O₃ stress is most consequential for soybean development; (iii) assess the potential for recovery following episodic O₃ exposure and how this differs between controlled and field conditions; and (iv) evaluate the capacity of remote sensing indicators, particularly SIF, to detect these processes at regional scale. By integrating experimental and remote sensing perspectives, our goal is to improve mechanistic understanding of O₃ stress in soybean and contribute to the development of remote sensing-based early warning systems and decision-support tools for farmers and agronomists.

## 2. Methodology

Our study integrates experiments and observations across spatial and temporal scales. We began with controlled, leaf-level measurements using a portable photosynthesis system (PPS) and extended the analysis to regional-scale using satellite remote sensing data. Conditions in the chamber experiment were designed to replicate the average temperature and relative humidity (RH) observed over four soybean growing seasons (2018-2021) in Crittenden County, Arkansas (Section 2.2) (**Figure S1**). The same environmental settings were applied during PPS measurements (Section 2.1) to ensure consistency across scales. This approach allowed us to compare chamber findings with multi-year satellite observations from TROPOMI and the Moderate Resolution Imaging Spectroradiometer (MODIS).

We emphasize that chamber and satellite results are not directly comparable because they represent very different spatial scales (leaf vs. field). Instead, our aim was to test whether the mechanisms of O₃ stress observed under controlled conditions also emerge at the regional scale, linking mechanistic understanding with applied monitoring. Based on the remote sensing data, soybean fields in Crittenden County did not experience numerous extreme weather or pollution events during the observed period (**Figures S1 and S2**), representing relatively healthy agroecosystems – making this study both more challenging and more relevant, as it examines O₃ effects under moderate, yet realistic, conditions.

### 2.1. Plant-Level Chamber Experiments

Soybeans (*Glycine max*) were grown from seed. Seeds were sown in small containers on 06 February 2025, and germination began on 12 February 2025. Plants grew in a homemade polytetrafluoroethylene growth chamber under controlled laboratory conditions with a standard 12-hour light–dark cycle. A Li-COR CO₂/H₂O gas analyzer (LI-840A) and a 2B-Technologies O₃ monitor (Model 202) continuously measured chamber CO₂ and O₃ mixing ratios. Other than during fumigation periods (described below), plants were exposed only to ambient laboratory O₃ levels, with daily means of ∼30 ppb.

The chamber (0.61 m × 0.61 m × 0.61 m) was made of transparent, chemical-resistant, polytetrafluoroethylene film with perfluoroalkoxy and polytetrafluoroethylene Swagelok fittings coupled with clear fluorinated ethylene propylene tubing (**Figure 1**). A 60 L food-grade CO_2_ cylinder (Sodastream) maintained mixing ratios in the chamber at ∼400 ppm. Lab air was pushed by a Gast compressor/vacuum pump (DOA-P704-AA) through a bubbler system with deionized water into the chamber to maintain RH at 65 ± 10% throughout the fumigation. Internal chamber temperature was 25 ± 2 °C. A 300 W LED grow light (MARS HYDRO) controlled light intensity at 1000 µmol m⁻² s⁻¹, measured at the top of the chamber.

**Figure 1.**
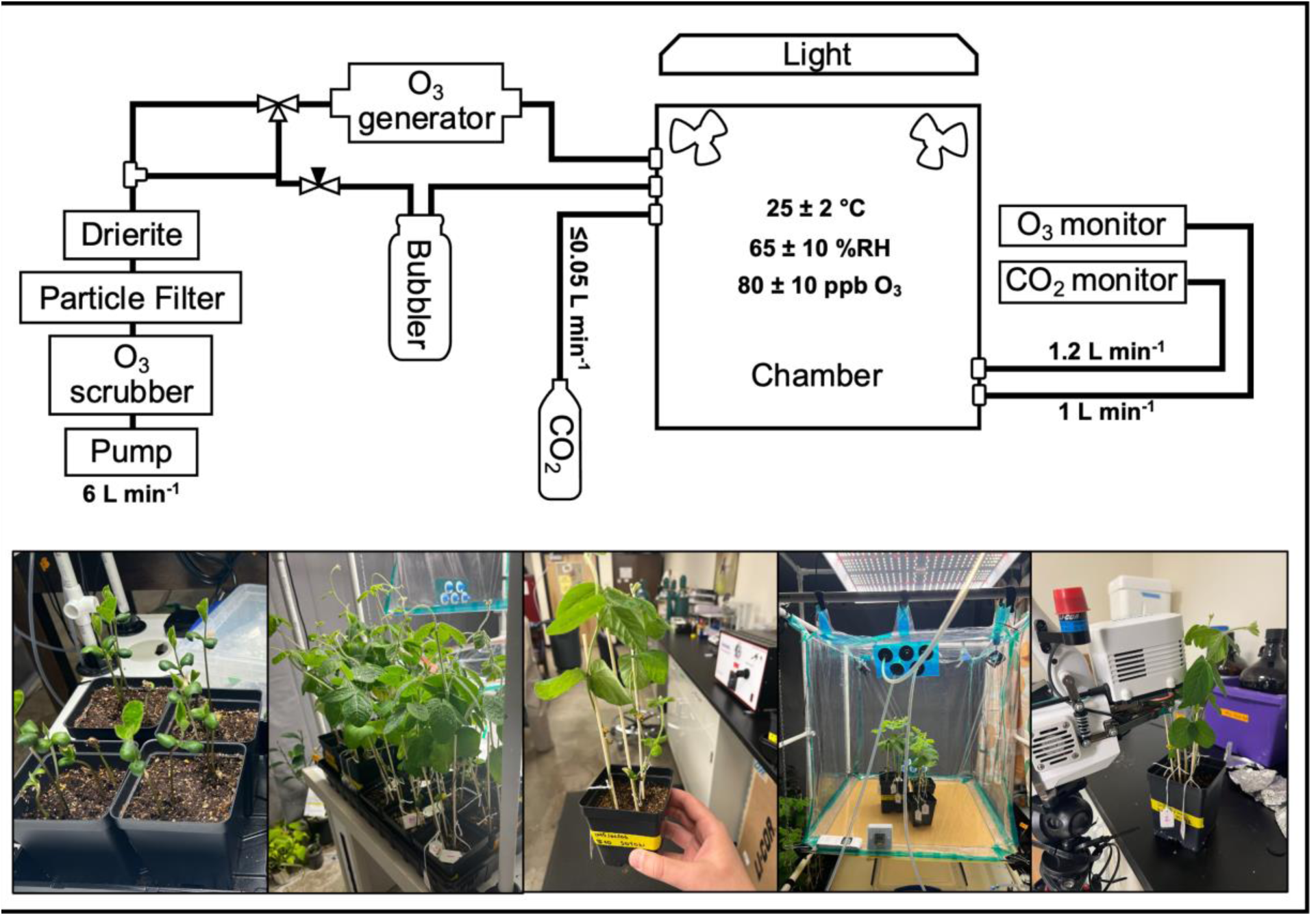
Schematic of the custom-built plant fumigation chamber (top) and photos from the soybean vegetative growth cycle during the experiment (bottom). Total inlet flow of 6 L min⁻¹ was split between the O₃ and humidification lines as needed to regulate RH and O₃.

The five-week plant-level experiment started on 03 March 2025. During this period, 1-3 leaves from each selected plant (five leaves in total) were measured weekly using a Li-Cor LI-6800 PPS. Parameters included CO₂ assimilation (A, rate of CO₂ movement through the stomata) stomatal conductance (Gs, movement of CO₂ and H₂O through the stomata) (Riches et al., 2020). We also measured maximum light-adapted fluorescence (Fm′) and steady-state fluorescence (Fs) which were used to calculate ΦPSII. PPS leaf chamber conditions were set to 26 ± 0.2 °C and 63 ± 0.5% RH, equivalent to 1.36 ± 0.04 kPa VPD. These conditions matched the growth chamber environment and were representative of field conditions in Crittenden County, Arkansas (see Section 2.2, **Figure S1**).

Plants were divided into two groups: three controls and three O₃ fumigation plants. Both groups followed the same measurement schedule and chamber conditions. Each week, fumigated plants were exposed for four hours to 80 ± 10 ppb O₃ generated by a pen-ray lamp (Analytik Jena AG) connected to the dry air inlet. PPS measurements were taken on five leaves immediately before each exposure (pre-treatment) following a 20-minute acclimation period, and again immediately after exposure (post-treatment). Control plants underwent the same procedure, but without O₃ fumigation. This cycle was repeated weekly for five consecutive weeks, during which plants advanced through vegetative growth and began pod development (**Figure 1**).

Being aware of the limited leaf-level dataset and the importance of distinguishing biological outliers from systematic ones (Alvarez Prado et al., 2019; Julián et al., 2024), we applied the stricter approach of Leys et al. (2013), excluding values outside three median absolute deviations (MADs) (**Figure S3**). This approach minimizes the impact of errors in measurements while maintaining the most representative physiological responses. We also note that measurements of A in week five may be confounded by plants nearing the end of their life cycle or by limitations to root growth from the small containers.

The results of the chamber experiment are presented in Section 3, including analyses of (i) soybean leaf responses to initial O₃ exposure, (ii) recovery potential between repeated exposures, (iii) effects of fumigation on plant growth, and (iv) the behavior of photosynthetic fluorescence parameters under subtle O₃ stress.

### 2.2. Remote sensing analysis

The chosen study area, shown on **Figure 2**, located in Crittenden County (Arkansas, U.S.), is a significant soybean-producing region in a state that ranks among the top soybean producers nationally (Ross, 2020). Due to the coarse spatial resolution of the TROPOMI SIF product (7 × 3.5 km), two points of interest were selected over a cluster of large soybean fields (AR1 and AR2).

**Figure 2.**
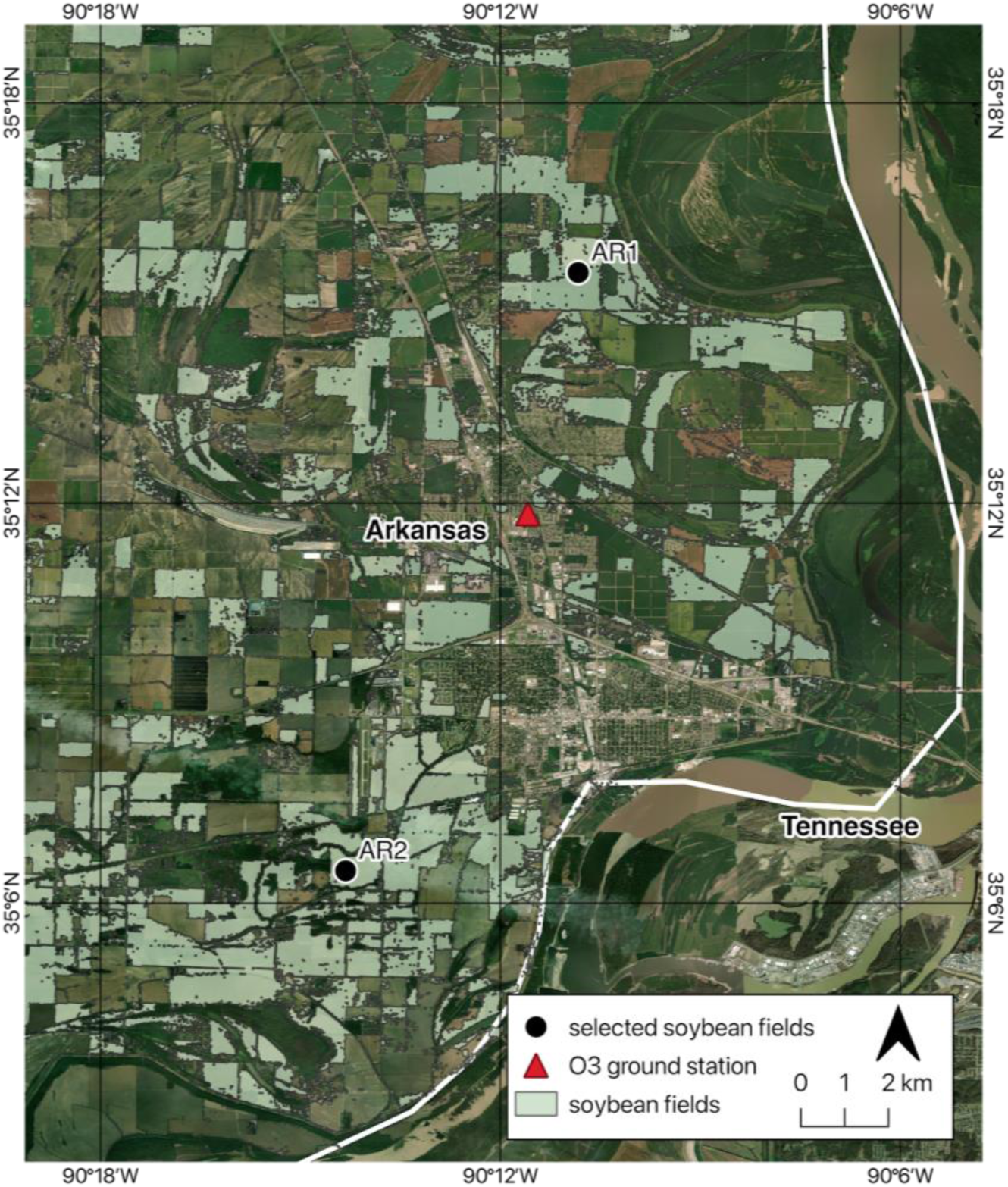
Study area in Crittenden County, Arkansas, U.S.

To evaluate the impact of O₃ on soybean fields at the regional scale, we integrated multiple datasets (**Table 1**) (**Figures 3 and S4**). Ground-level O_3_ data from a nearby Environmental Protection Agency (EPA) monitoring station in Crittenden County provided daily 8-hour means and hourly concentrations, the latter used to calculate weekly AOT40 by summing exceedances of O_3_ above 40 ppb during daylight hours (08:00-20:00). Daily meteorological variables, including maximum temperature (T_max_), RH, VPD, grass reference evapotranspiration (ETo), and precipitation (PREC) were obtained from the University of Idaho GRIDMET dataset (Abatzoglou, 2013). MODIS datasets provided vegetation and productivity indicators, including 8-day cumulative gross primary production (GPP, the total carbon fixed by plants through photosynthesis summed over 8-day period) (Robinson et al., 2018), 4-day composite fraction of photosynthetically active radiation (fPAR, the portion of incoming sunlight absorbed by the canopy), 4-day composite leaf area index (LAI, one-sided green leaf area per unit ground area), daily NDVI (derived from surface reflectance, as a measure of canopy greenness), and 3-hourly total photosynthetically active radiation (PAR, incident solar radiation in the visible spectrum (400-700 nanometers)).

**Figure 3.**
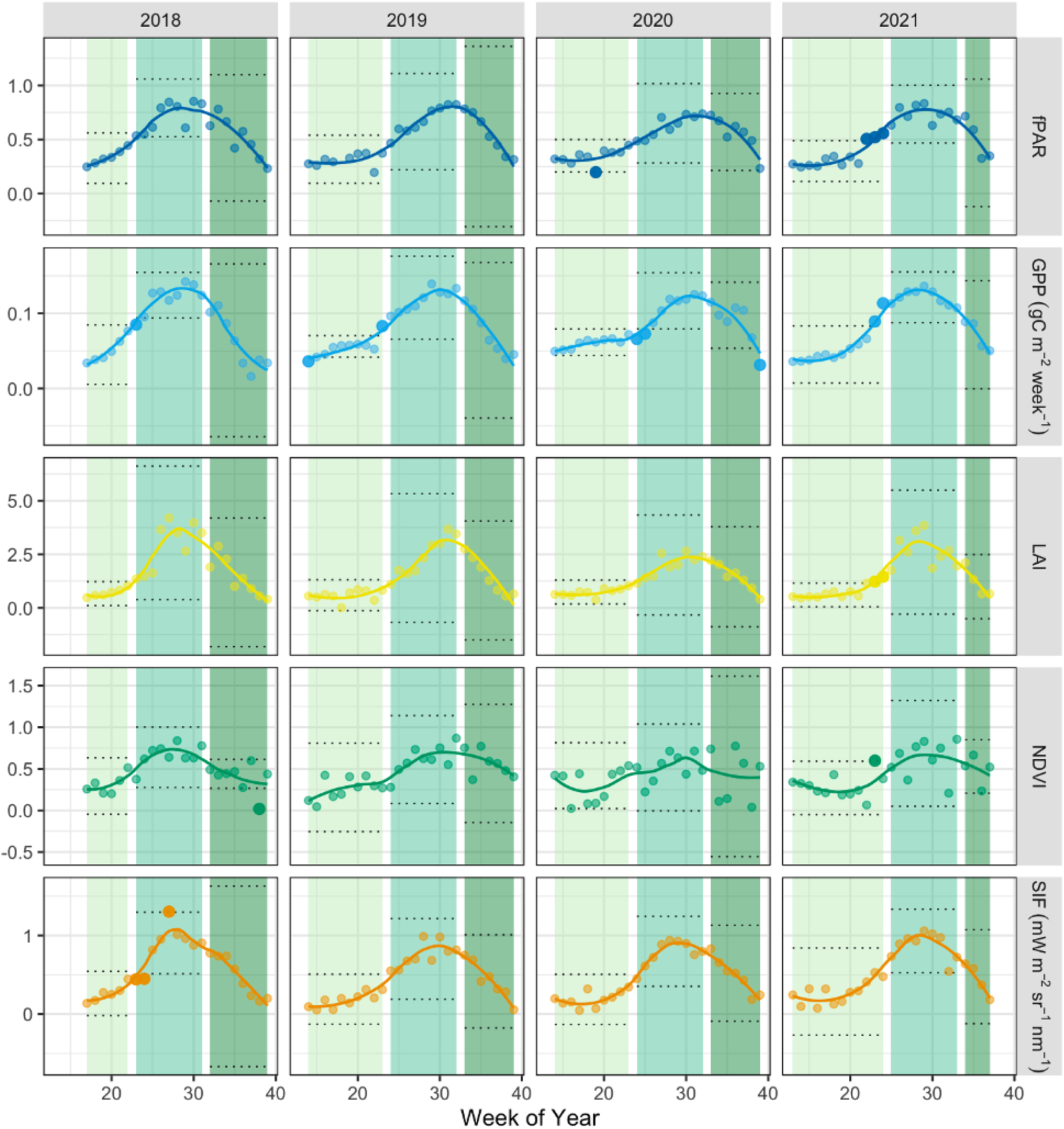
Weekly time series of vegetation indicators (fPAR, GPP, LAI, NDVI, and SIF) in soybean fields in Crittenden County, Arkansas, across four growing seasons (April-September, 2018-2021). Points show weekly observations, and lines represent smoothed seasonal signatures. Shaded bands indicate the early (light green), peak (mid green), and late (dark green) phases of the growing season. Solid (non-transparent) points mark flagged outliers within each phase.

**Table 1.** Specifications of datasets used in this study for remote sensing analysis.

| Data | Description | Units | Temporal resolution | Spatial resolution | Source |
| --- | --- | --- | --- | --- | --- |
| O<sub>3</sub> | In-situ ozone | ppb | Daily | / | EPA |
| SIF | Solar-induced fluorescence at 743 nm | mW m<sup>-2</sup> sr<sup>-1</sup> nm<sup>-1</sup> | Daily | 7 × 3.5 km | ESA-TROPOSIF |
| T<sub>max</sub> | Maximum temperature | °C | Daily | 4.6 km | GRIDMET |
| RH | Relative humidity | % | Daily | 4.6 km | GRIDMET |
| VPD | Vapor pressure deficit | kPa | Daily | 4.6 km | GRIDMET |
| PREC | Precipitation amount | mm | Daily | 4.6 km | GRIDMET |
| ET<sub>o</sub> | Grass reference evapotranspiration | mm | Daily | 4.6 km | GRIDMET |
| GPP | Gross primary production | kgC m<sup>-2</sup> 8-day<sup>-1</sup> | 8-day | 0.3 km | MODIS |
| PAR | Photosynthetically active radiation | W m<sup>-2</sup> | 3-hour | 0.5 km | MODIS |
| fPAR | Fraction of photosynthetically active radiation | / | 4-day | 0.5 km | MODIS |
| LAI | Leaf area index | / | 4-day | 0.5 km | MODIS |
| NDVI | Normalized difference vegetation index | / | Daily | 0.5 km | MODIS |

SIF 743 nm data were derived from the ESA-TROPOSIF project (Guanter et al., 2021), which uses daily Sentinel-5P TROPOMI measurements. Data from May 2018 to December 2021 were extracted over selected points, filtered to remove negative and extreme values, and averaged between fields to maximize signal coherence. The analysis period covered four soybean growing seasons, each defined as the period from April to September. Within each season, we further subdivided the timeframe into early, peak, and late stages based on SIF dynamics. The peak season range was defined as the four weeks before and after the maximum SIF value. The four weeks prior to the peak period were defined as the early season, and the four weeks after the peak period were defined as the late season, corresponding with senescence and harvesting (**Figure 3**).

We aggregated all datasets to a weekly scale for analysis. Weekly water balance (WB) was calculated as the difference between accumulated PREC and ETo. Each of the eight 3-hourly PAR bands per day (in W m⁻²) was converted to photons using the standard factor for sunlight (1 W m⁻² ≈ 4.57 µmol m⁻² s⁻¹) and integrated over 10 800 s (3 hours), summed across the day and then across seven days to obtain weekly incident PAR (mol m⁻² week⁻¹) (Lozano et al., 2023; Wang et al., 2020). 8-day GPP was first divided by 8 to get approximate daily values and then summed across week. We calculated weekly absorbed photosynthetically active radiation (APAR) as APAR = PAR × fPAR, which was used to calculate weekly light use deficiency (LUE) as LUE = GPP/APAR, and SIF_yield_ = SIF/APAR (efficiency of energy conversion from absorbed light into fluorescence) (Magney et al., 2019; Miao et al., 2018). Following the framework of (Massmann et al., 2019), we further estimated Gs as a function of GPP and VPD (E1):

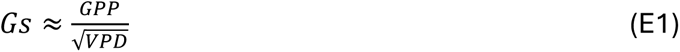

This proxy (E1) assumes that plants open their stomata during photosynthesis (higher GPP) and close them when the atmosphere becomes drier (higher VPD), helping estimate potential O₃ uptake.

The remote sensing allowed us to assess O₃ impacts on soybean physiology at the regional scale across four growing seasons. In Section 4, we present these results, including analyses of (i) the effects of early versus late O₃ episodes, (ii) the potential for recovery in the weeks following exposure, (iii) the relationship between seasonal O₃ accumulation and yield proxies, and (iv) changes in the coupling between vegetation health indicators under different O₃ conditions. Comparisons with plant-level chamber experiments are made throughout the section to evaluate whether the mechanisms observed at the leaf level can also be detected across larger spatial and temporal scales.

## 3. Plant-level O_3_ Fumigation Experiment

### 3.1. Impact of initial O_3_ exposure on photosynthesis

Initial exposure (Week 1) to O_3_ had the most pronounced immediate effect on all measured physiological parameters (**Figure 4**). In the first week of fumigation, treated plants showed a stronger decrease in the ratio of post- to pre-chamber values for both A and Gs compared to controls, indicating early and acute stress response to O_3_. This immediate response, which was not observed in following exposures, suggests that plants were not acclimated to the elevated O_3_ and had not yet activated defense mechanisms, consistent with earlier reports that initial O_3_ events trigger strong physiological reactions in soybean (Morgan et al., 2003; Singh et al., 2009). These findings highlight the acute sensitivity of soybean plants during the early vegetative stage and support the idea that timing of exposure is as important as intensity. That is, elevated O₃ early in the growing season is likely to cause more lasting damage on photosynthesis than equivalent exposure later.

**Figure 4.**
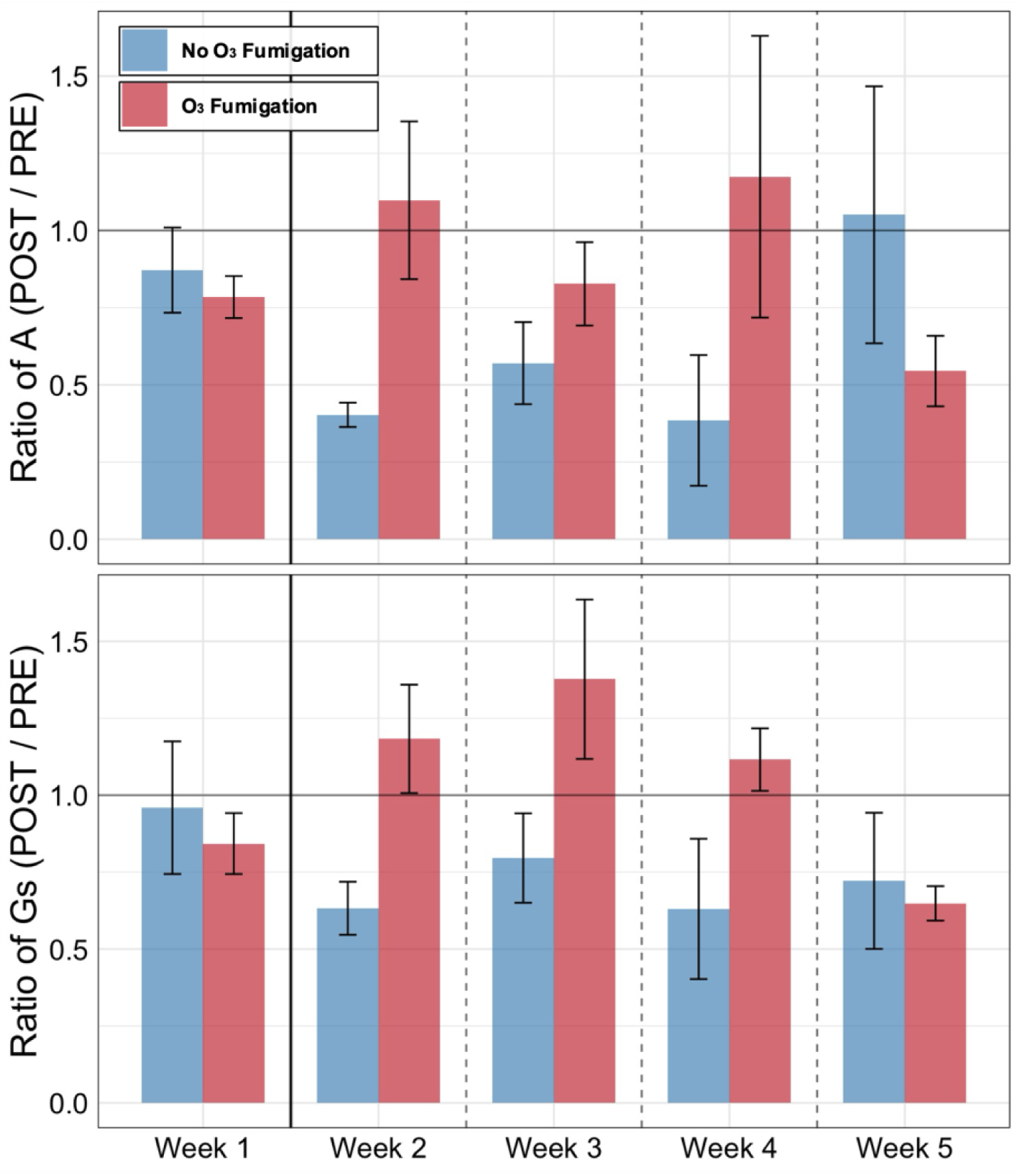
Physiological responses of soybean plants to O_3_ fumigation over five weeks are expressed as the ratio of the post- and pre-chamber measurements of A (top row), and Gs (bottom row). Bars represent the mean ratio (n = 5 leaves) ± standard error (SE) for each week of exposure for O₃-treated (red) and control (blue) plants.

In the following weeks, the pattern was reversed: control plants often showed stronger negative post-chamber reactions than fumigated plants. This finding suggests that the chamber environment itself is a stressor, confounding immediate post-exposure responses and complicating the separation of O₃ effects from chamber effects. Adding O₃ as an additional stressor may have altered or masked immediate post-exposure signals, leading to inconsistent ratios. For this reason, we interpret pre-chamber measurements in subsequent weeks as more reliable indicators of long-term O₃ impact, revealing the potential for recovery, which we explore in the next section.

### 3.2. O₃ exposure limits plants’ ability to recover photosynthesis and efficiency

Focusing only on pre-chamber values allows us to assess how much plants recovered between exposures. Soybean plants showed limited resilience in photosynthetic function (**Figure 5**). After the first fumigation, A and Gs never fully returned to pre-exposure levels in O₃-treated plants – in clear contrast to control plants, which maintained stable or increasing values across the same period. Similarly, ΦPSII declined and remained suppressed in fumigated plants, as opposed to controls. This trend indicates that O₃ not only limits carbon assimilation and gas exchange, but also impairs photochemical efficiency, further constraining recovery potential. Thus, while the impact of the first O₃ treatment may not have been extreme relative to the control on immediate, post-treatment parameters, plant damage was clear within a week, reflecting interrupted development from the modest four-hour O₃ fumigation at 80 ppb. Thus, O₃ exposure not only causes an immediate – albeit subtle – stress on the time scale of hours, but also causes a longer-term response of suppressed growth on the time scale of days to weeks. These findings align with other controlled studies showing that repeated O₃ exposures compromise recovery processes such as stomatal reopening and enzymatic function in the Calvin cycle (Morgan et al., 2003; Osborne et al., 2016). Suppression of ΦPSII is consistent with reports of O₃-induced damage at the chloroplast level (Chen et al., 2009), as well as with declines in canopy-scale fluorescence under elevated O₃ (Wu et al., 2024).

**Figure 5.**
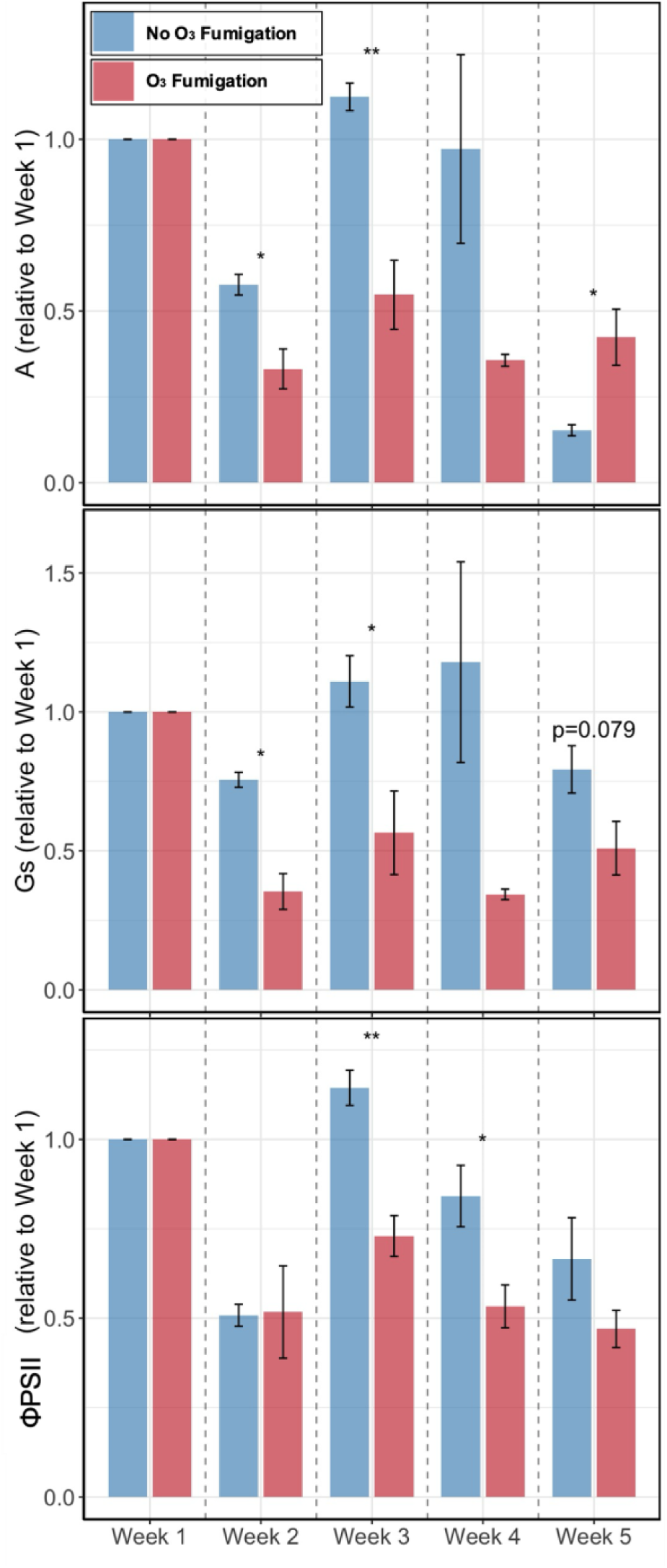
Pre-chamber measurements of A (top), Gs (middle) and ΦPSII (bottom) relative to week 1, in O_3_-treated (red) and control (blue) soybean plants across five consecutive weeks. Bars are mean values ± SE (n = 5 leaves). Asterisks indicate significance levels (* p < 0.05, ** p < 0.01).

Pre-chamber measurements show that O₃-treated plants failed to recover photosynthetic function between exposures, with sustained suppression of A, Gs, and ΦPSII indicating an ongoing physiological burden that extends beyond short-term inhibition and is reflected in the slower growth patterns described in the following section.

### 3.3. Subtle O₃ exposure suppresses soybean growth and biomass

O_3_ exposure impacted morphological development. Weekly measurements showed stalk height increasing more slowly in O_3_-treated plants than in control plants (**Figure 6**). By the end of the five-week experiment, average plant height in the fumigated group was consistently lower, even though no visible foliar damage was apparent. Final dry weight (DW) measurements supported this pattern, with non-fumigated plants accumulating more biomass (0.26 ± 0.04 g DW) than fumigated plants (0.21 ± 0.03 g DW). This growth suppression under ambient O_3_ aligns with prior findings that O₃ reduces biomass accumulation even in the absence of necrotic symptoms (Morgan et al., 2003). This effect is likely caused by reduced carbon uptake due to the observed lower assimilation rates (see Section 3.2), combined with a shift in resource allocation from growth processes to antioxidant defense mechanisms (Osborne et al., 2016).

**Figure 6.**
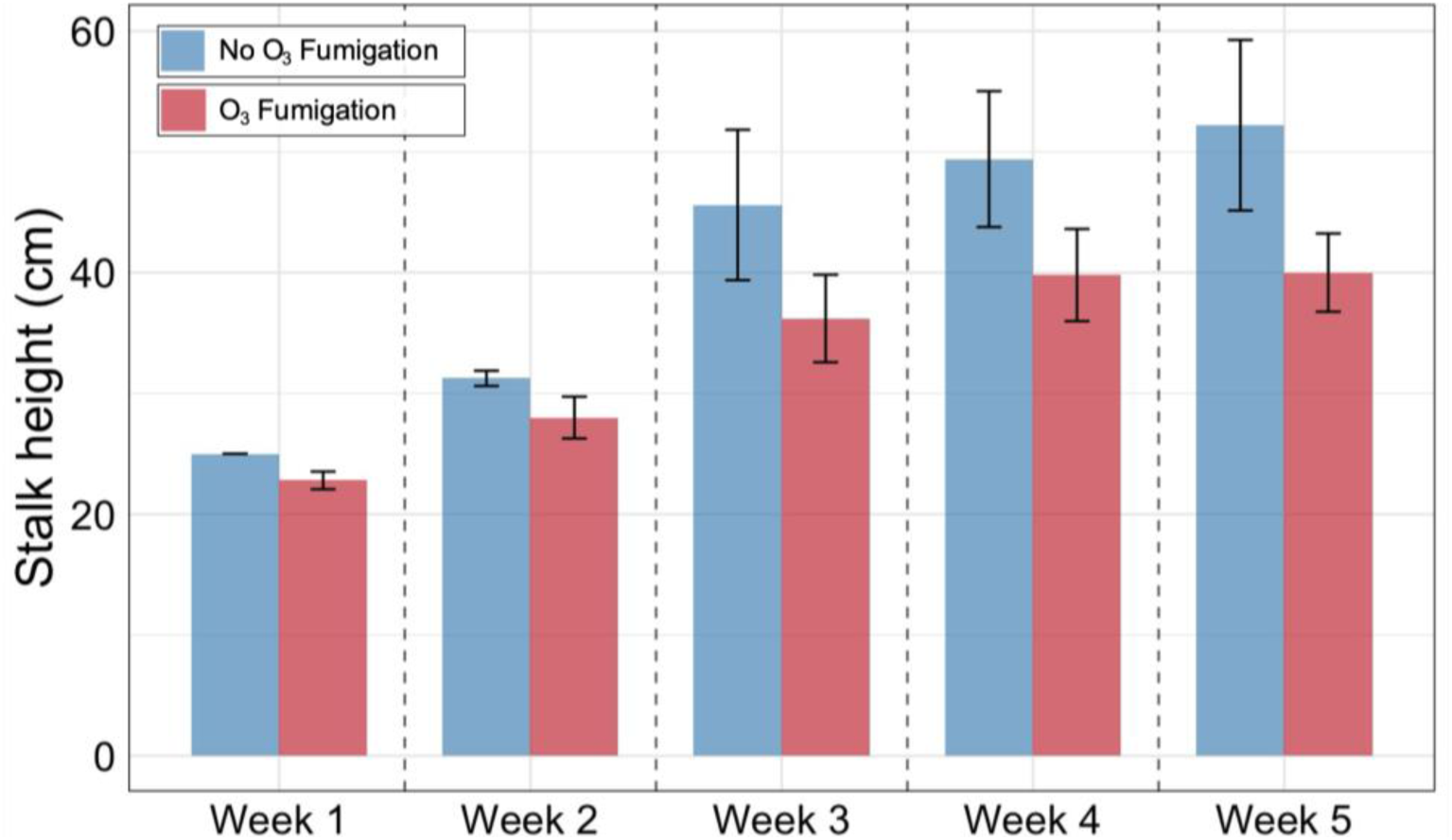
Weekly stalk height measurements of soybean plants under O₃ fumigation (red) and control conditions (blue) over a five-week period. Bars represent mean stalk height ± SE (n = 5 stalks).

The impact of fumigation on growth persisted despite some recovery of gas exchange parameters in later weeks (see Section 3.2), suggesting that the effect of O₃ on developmental processes extends beyond short-term photosynthetic inhibition. This reinforces the concept that chronic, sub-lethal O₃ exposure slows physiological development, especially when plants are exposed early in their growth cycle. These results are consistent with SoyFACE observations, where long-term O_3_ exposure reduced canopy height and biomass even when visual damage was absent (Ainsworth et al., 2014; Wu et al., 2024). However, our results further demonstrate that even more subtle and occasional changes in O₃, consistent with realistic scenarios currently occurring in agricultural regions, are sufficient to suppress plant growth and reduce biomass.

### 3.4. Long-term O₃ exposure decouples photosynthetic efficiency from fluorescence signals

Further analysis of pre-chamber values, discussed in Section 3.2, revealed a decoupling of ΦPSII from both, Fm′ and Fs, under O₃ stress (**Figure 7**). ΦPSII, which is calculated as (Fm′ - Fs)/Fm′, is usually tightly linked to these fluorescence components, reflecting a balance between absorbed light energy, photochemistry, and non-photochemical quenching (Oxborough & Baker, 1997).

**Figure 7.**
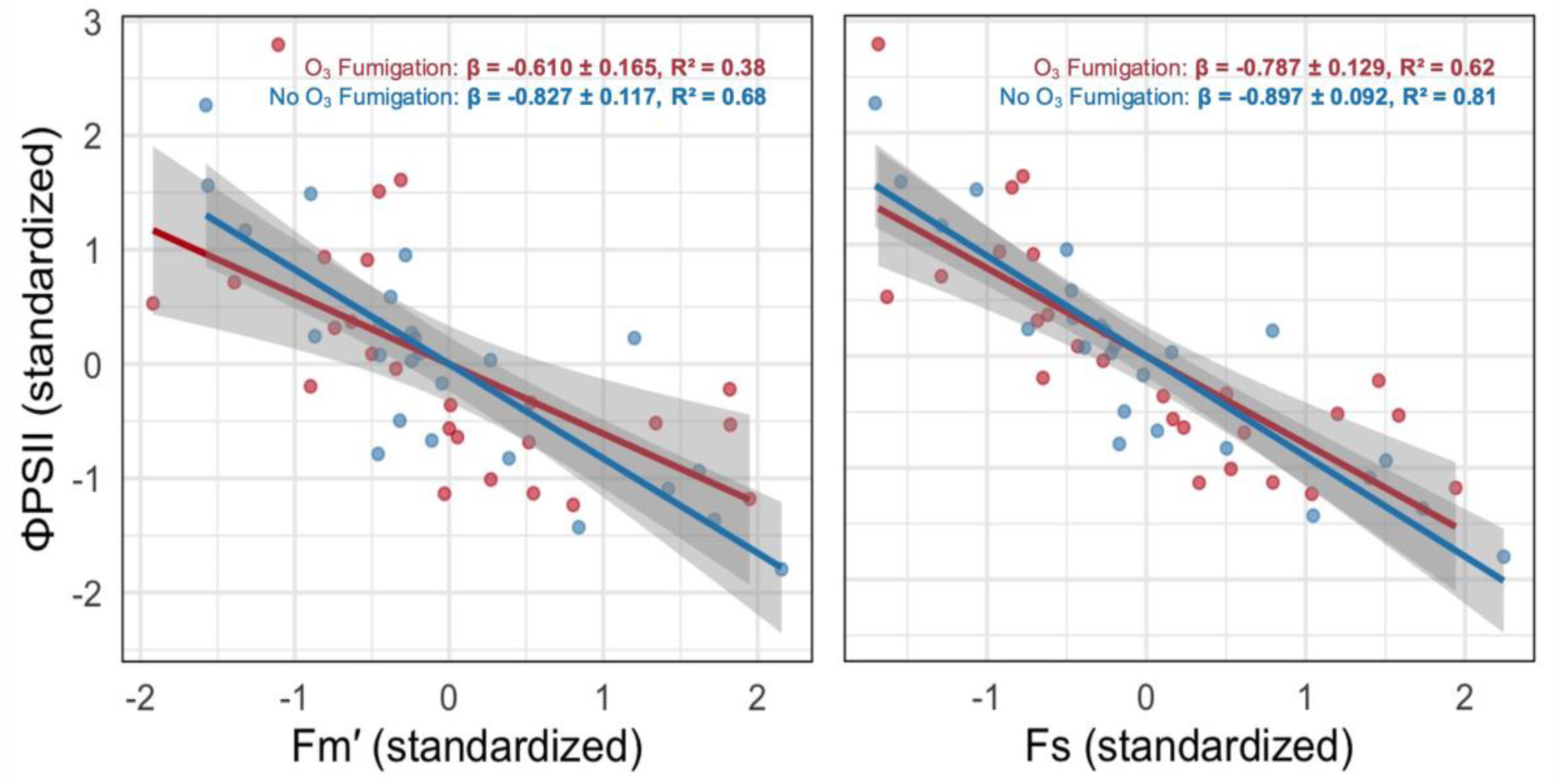
Relationships between ΦPSII and fluorescence components (Fm′ - left, and Fs - right) under O₃ fumigation (red) and control conditions (blue) presented by pre-chamber measurements across five weeks. Slopes (β ± SE) and R² are shown for each treatment; p- values < 0.001.

Observed decoupling indicates an imbalance between electron transport and energy dissipation, characteristic of strain-phase stress responses (Meroni et al., 2009), which implies that plants under chronic O₃ exposure may dissipate light energy in ways that no longer track with photochemical efficiency, an early-warning signal of impaired photosynthetic regulation (Ruban & Murchie, 2012). Furthermore, Betzelberger et al. (2012) argue that ΦPSII is among the most sensitive fluorescence-derived parameters to O₃ stress in soybean, even when gas exchange traits partially recover and that indices based solely on Fs may fail to detect strain, highlighting why ΦPSII offers a more mechanistic indicator of early stress.

Note that discussed decoupling was not observed in immediate post-chamber measurements, where Fm′ and Fs maintained similar slopes and R² with ΦPSII for fumigated and non-fumigated plants (**Figure S5**). This close relationship reinforces the idea, emphasized in previous sections, that the impact of O₃ is not immediately visible after exposure, but emerges over longer timescales and disrupts the functional link between fluorescence signals and ΦPSII efficiency.

## 4. Remote Sensing Analysis

### 4.1. Effect of first O_3_ episode in the growing season is more visible than equivalent exposure later

To test whether the timing of O₃ stress modulates crop responses, we identified O₃ episodes as weeks when AOT40 increased by more than 350 ppb·h relative to the previous week. For each of the four growing seasons (2018-2021), the earliest such episode was paired with a later episode of equivalent magnitude within the same season (n = 4, **Table S1**). To minimize influence of other environmental variables, the later episode was chosen to match first one to have similar PAR, WB, RH, VPD, and T_max_ values. Remote sensing vegetation indicators were compared across first versus later O₃ episodes (**Figure 8**). Another approach to minimizing seasonality trends was residualizing vegetation indicators against same meteorological covariates; results from the two approaches were consistent (**Figure S6**).

**Figure 8.**
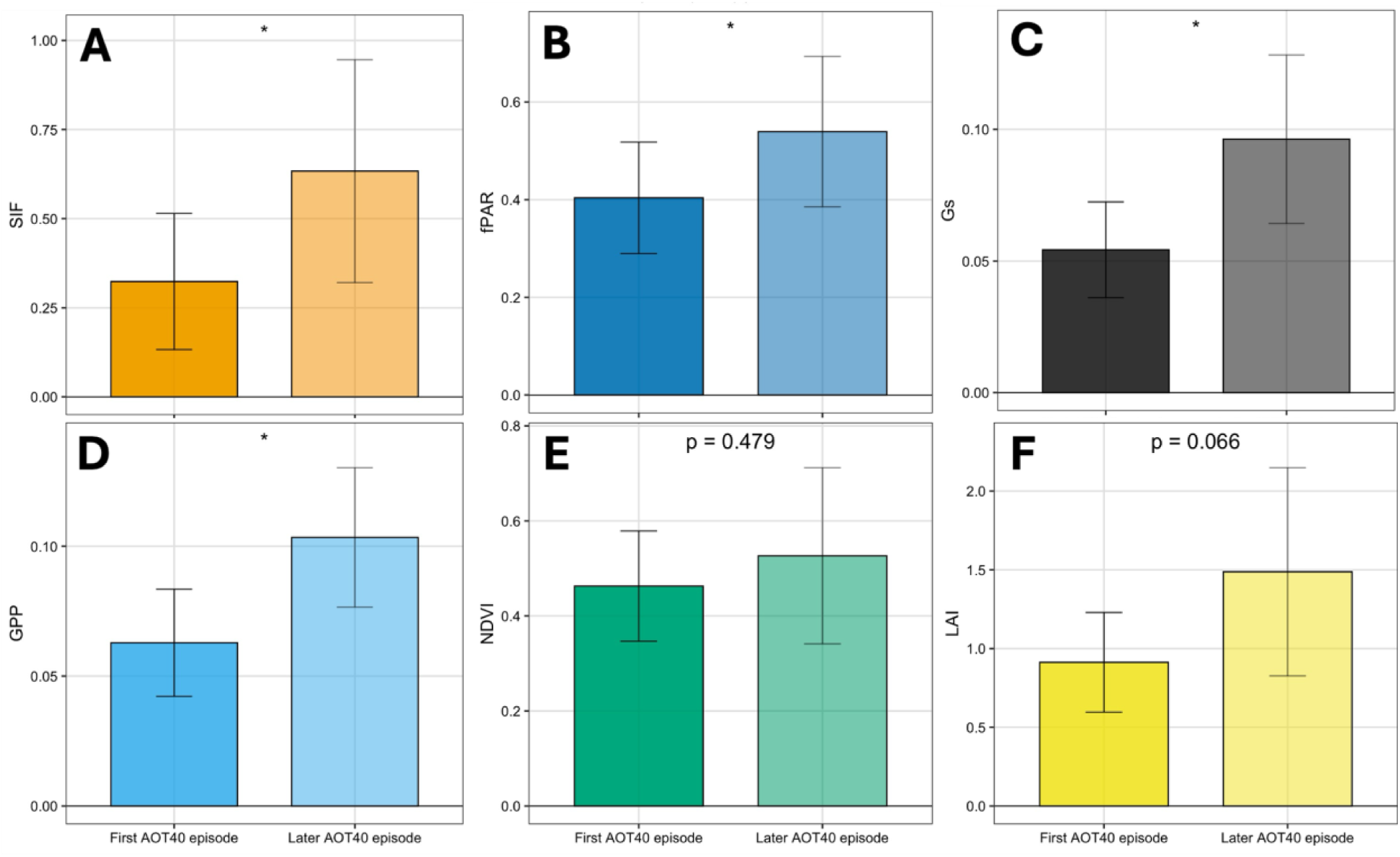
Vegetation health indicators values compared between the first and later O₃ episodes of equivalent AOT40 increase (>350 ppb·h) within each growing season (2018-2021). Bars represent mean ± SE across four paired episodes. Asterisks indicate significance levels (* p < 0.05).

Across all four years, the first O₃ episode of the season had a significantly stronger negative impact on functional indicators compared to equivalent exposures later in the season. SIF, GPP, fPAR, and Gs all showed significantly lower values during early-season episodes, while NDVI and LAI, which follow structural canopy development were not significantly different between two episodes. These findings mirror the leaf-level results (Section 3.1), where the first O₃ fumigation caused the largest drop in A and Gs, while following exposures triggered more muted responses as plants initiated partial defense mechanisms. Together, our results emphasize that the soybean is particularly vulnerable to O_3_ early in its vegetative cycle, when photosynthetic capacity and resource allocation patterns are still being established (Betzelberger et al., 2012).

Physiologically, early vulnerability could be the result of the rapid development of sink capacity, which promotes O₃ uptake and amplifies oxidative stress (Morgan et al., 2003; Singh et al., 2009). At this stage, antioxidant defenses and photoprotective processes have not yet fully developed, leaving ΦPSII particularly vulnerable to disruption (Chen et al., 2009; Osborne et al., 2016). This seasonal sensitivity is consistent with FACE experiments, in which chronic O₃ exposure accelerated senescence and reduced photosynthetic capacity, particularly during reproductive development (Ainsworth et al., 2014; Wu et al., 2024). Our results extend these insights by demonstrating that short-term, episodic O₃ events, found in agricultural regions – well below regulatory thresholds and chronic FACE exposures – can induce measurable strain when they occur early in the season. The strong suppression of SIF (A) and GPP (D) during these episodes underscores their value as functional indicators for detecting O₃ stress before visible canopy changes occur (E and F), in line with regional studies reporting stronger SIF-O₃ coupling during early vegetative stages (Feng & Kobayashi, 2009; Montes et al., 2022).

### 4.2. Real-world O₃ episodes slow down plant recovery

Using the O₃ episodes identified in Section 4.1 (ΔAOT40 > 350 ppb·h), we observe how soybean vegetation health indicators recover from their first seasonal O₃ event compared to a later event (**Figure 9**). We are aware that values of our vegetation indices could in principle be influenced by differences in canopy development between early and later episodes. However, as shown in Section 4.1, structural parameters such as NDVI and LAI did not differ significantly between the paired events, suggesting that canopy development was not a major confounding factor.

**Figure 9.**
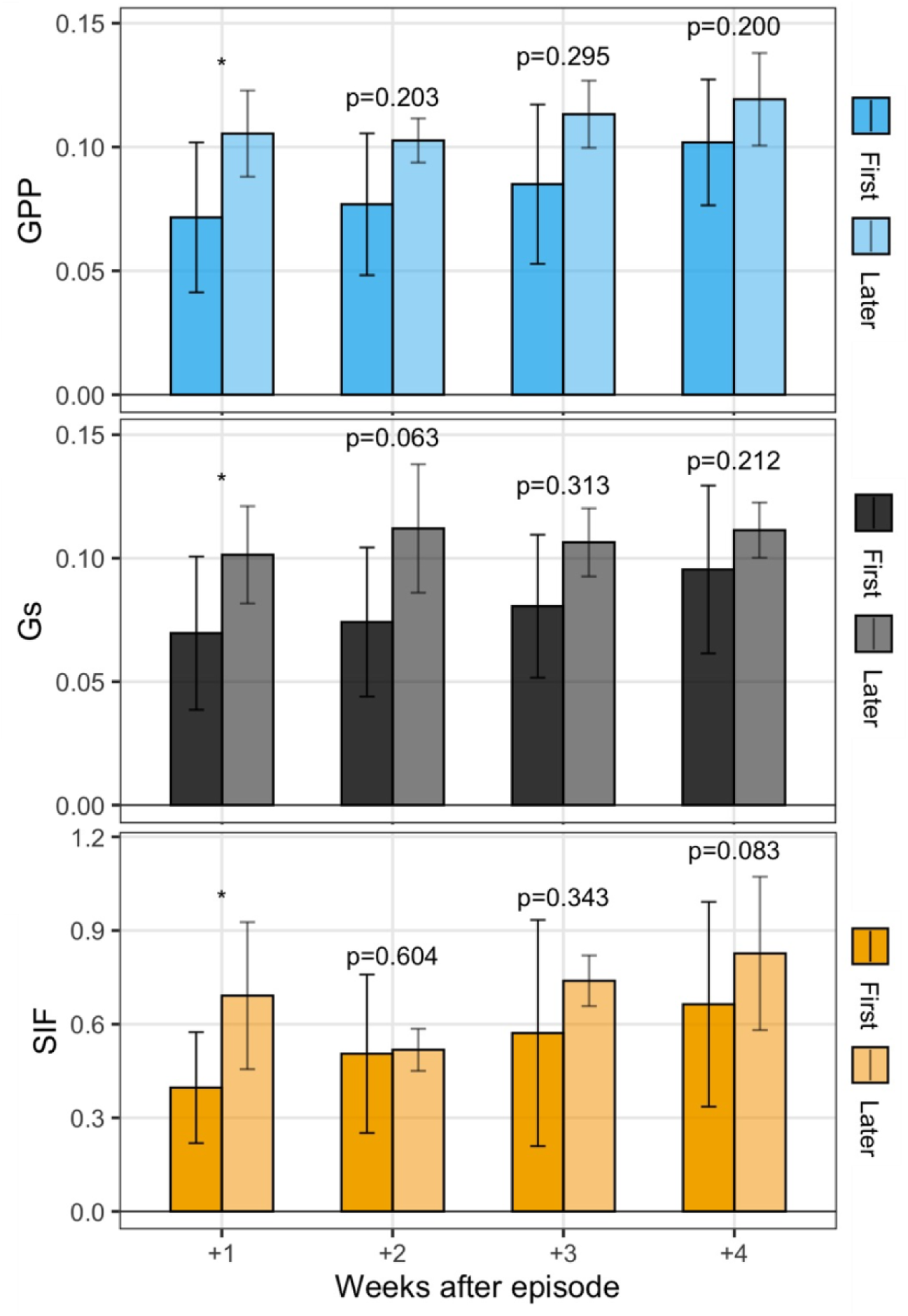
Recovery of functional vegetation indicators (GPP, Gs, and SIF) following the first (dark bars) vs. later (light bars) seasonal O₃ episodes (ΔAOT40 > 350 ppb·h). Bars show mean ± SE across four growing seasons (2018-2021). Asterisks indicate significance levels (* p < 0.05).

Results show that functional indicators (GPP, Gs, and SIF) were consistently more suppressed in the week following the first seasonal O₃ episode, compared to later episodes. In subsequent weeks, the gap between early and later episodes narrowed, suggesting canopy-scale recovery. This recovery contrasts with the chamber experiment (Section 3.2), where fumigated plants failed to recover fully between weekly exposures. The consistent stability of GPP, Gs, and SIF after later-season O₃ episodes suggests that plants may adapt to previously experienced strain. Structural indicators such as LAI, fPAR, and APAR (**Figure S7**) showed similar patterns – significant suppression after early episodes followed by recovery. Notably, LAI indicates that canopy development slowed after the first O₃ episode, compared to later ones, consistent with chamber findings on growth suppression (Section 3.3).

The multi-week recovery lag observed after early episodes aligns with leaf-level evidence from Section 3.2. As the season progresses, acclimation and phenological shifts likely shorten this lag by increasing sink strength, enhancing antioxidant defences, and adjusting stomatal sensitivity, buffering transient O₃ pulses (Betzelberger et al., 2012; Montes et al., 2022; Osborne et al., 2016). These processes, and acclimation to an earlier strain phase (Meroni et al., 2009), may explain the reduced sensitivity to later O₃ episodes.

### 4.3. Higher O₃ accumulation in early and peak growing season reduces final soybean yield

We used cumulative GPP as a proxy for soybean yield (Dold et al., 2019; Marshall et al., 2018; Wurster et al., 2021) to examine how seasonal accumulations of AOT40 translate into productivity losses (**Figure 10**). Results show that total accumulated O₃ across the full season did not correlate with yield (R² = 0.04, p = 0.81). Instead, exposure during the early growing season alone explained 47% of variance in cumulative GPP, and when early and peak seasons were combined, the relationship strengthened dramatically (R² = 0.98, p = 0.01). The strong predictive power of early and peak season exposure for final yield also aligns with meta-analyses showing that reproductive and vegetative transitions are the most sensitive phases for soybean yield determination (Feng & Kobayashi, 2009; Montes et al., 2022).

**Figure 10.**
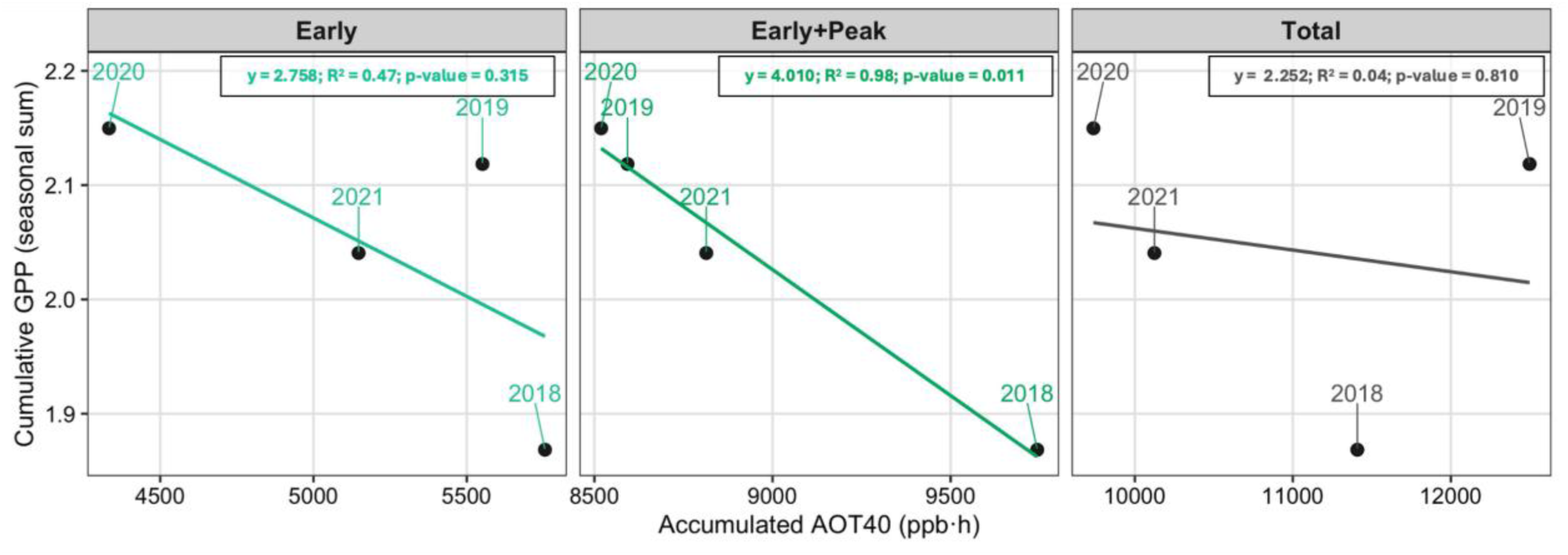
Seasonal cumulative GPP as a proxy for soybean yield plotted against accumulated AOT40 (ppb·h). Panels show relationships for early season only (left), early + peak seasons (middle), and the total growing season (right). Lines indicate linear regression fits with corresponding R² and p-values.

This finding reinforces our plant-level results (Section 3.3), where growth suppression occurred even in the absence of visible foliar damage. Furthermore, we connect findings from Sections 3.1 and 4.1, which showed that early-season exposures leave lasting impacts while later episodes have weaker effects, highlighting that it is not the total seasonal O₃ dose that determines yield losses, but rather when an elevated exposure event occurs. Early and peak stages, when plants are expanding canopies and establishing reproductive sinks, are the most consequential for O₃ impacts (Ainsworth et al., 2014; Morgan et al., 2003).

For real-world application, we show that monitoring and mitigating O₃ during the early and peak growing season is critical for protecting yields, shedding light on potential mechanisms behind regional yield losses reported in the U.S. and Asia, where seasonal O₃ episodes often cluster in late spring and early summer (Leung et al., 2022; Tai et al., 2021). Early strain detection using SIF and other functional indicators, as demonstrated in previous sections, could allow agronomists and farmers to take adaptive actions before the damage phase occurs and losses become irreversible. The seasonal distribution of O₃ further supports this conclusion. For example, 2019 had the highest total accumulated AOT40 but yielded relatively well because much of the exposure occurred late in the season (**Figure S8**). By contrast, the high combined early and peak season exposure of 2018 had the lowest yield. These results emphasize that policies and crop management strategies must consider seasonal timing of O₃ exposure, not just seasonal totals or regulatory exceedances.

### 4.4. Accumulated O₃ affects relationships between remote sensing vegetation health indicators

We split the early and peak dataset into AOT40 tertiles (≤311, 311-508, ≥508 ppb·h) and, within each class, fit standardized regressions. We define these tertiles as ‘lower’, ‘middle’, and ‘higher’ O_3_ exposure. Relationships between GPP-SIF, Gs-SIF, LAI-SIF, fPAR-GPP, and fPAR-SIF, stayed statistically strong across all classes (**Figure S9)**; the main changes were modest shifts in explained variance rather than large slope breaks. The fact that all of the relationships remained significant across classes suggests that basic canopy coordination is maintained, even if stress reduces its strength.

Moving beyond GPP-SIF-fPAR-Gs relationships, we also examined how O₃ alters broader coordination between light absorption, fluorescence, and efficiency. **Figure 11** shows six additional relationships (APAR-GPP, APAR-SIF, LUE-Gs, LUE-SIF, SIFyield-Gs, and SIFyield-LUE) across AOT40 tertiles. These metrics explicitly link absorbed energy, stomatal control, and realized carbon gain, and therefore allow us to assess whether O₃ disrupts not just single pairings but the broader chain of processes from light capture to photosynthesis.

**Figure 11.**
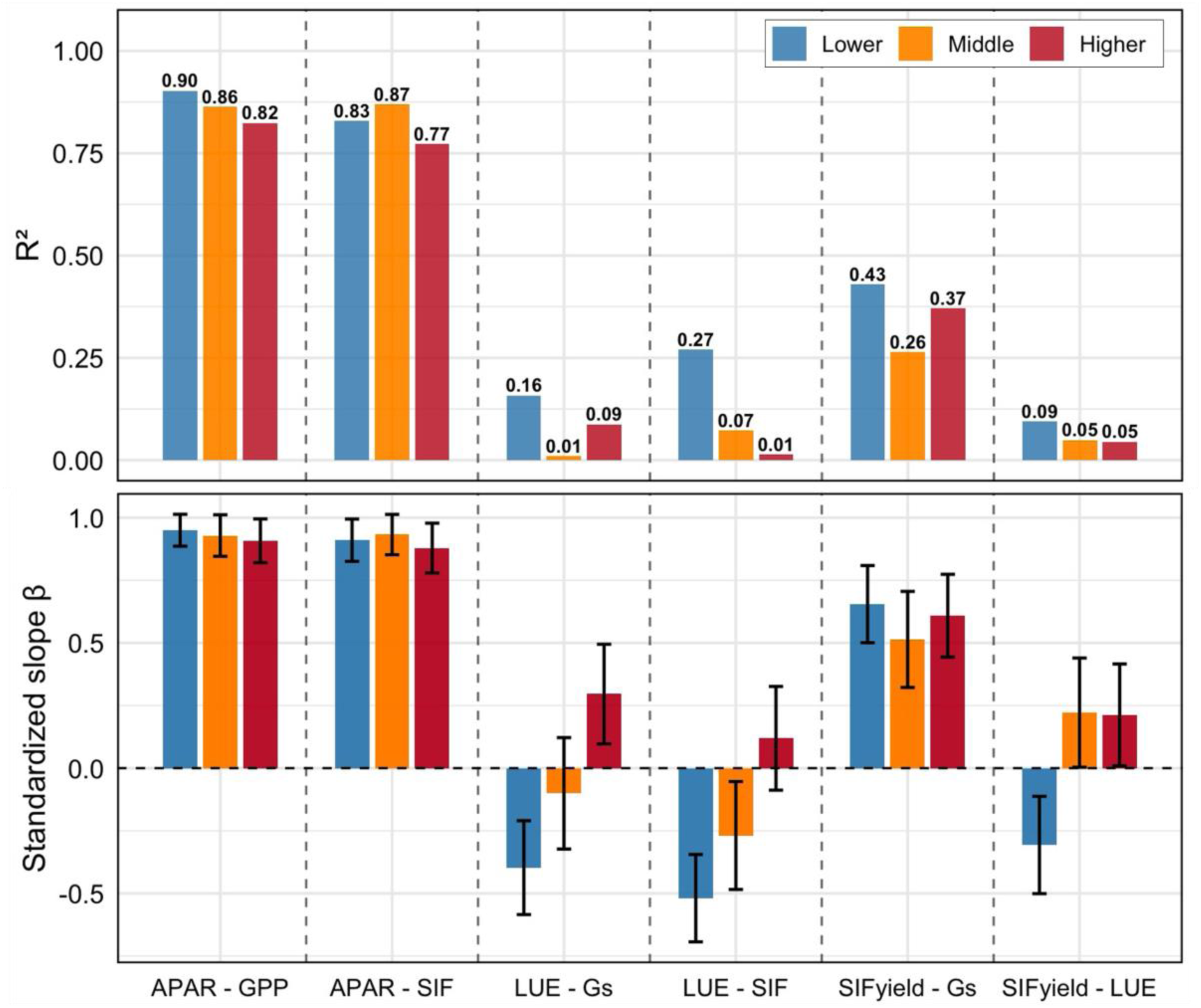
Relationships between vegetation health indicators during the early and peak soybean growing season by different O_3_ exposure. Bars show standardized regression slopes (β, bottom) ± SE, and coefficients of determination (R², top) for pairs of functional and structural indicators. Lower (blue), middle (orange), and higher (red) tertiles correspond to AOT40 thresholds of ≤311, 311-508, and ≥508 ppb·h, respectively.

Relationships linking absorbed light to productivity (APAR-GPP) and fluorescence (APAR-SIF) remained strong across all classes but weakened under high O₃ which indicates that while canopies continued absorbing and re-emitting light, their ability to convert light into carbon gain declined, reflecting reduced LUE (Betzelberger et al., 2012; Osborne et al., 2016). Similarly, the already weak LUE-Gs relationship eroded with higher O₃, consistent with additional biochemical limitations beyond stomatal control (Clifton et al., 2020).

Relationships between efficiency metrics collapsed at high O₃. The LUE-SIF correlation, low, but present under lower exposures, disappeared in the highest class, suggesting that fluorescence no longer tracks photosynthetic efficiency under O_3_ stress. This canopy-scale pattern mirrors the leaf-level decoupling observed between ΦPSII and Fm′/Fs in Section 3.4, reinforcing that O₃ disrupts the coherence between photochemical signals and productivity across scales (Chen et al., 2025). On the other hand, SIFyield-LUE was almost zero under all O₃ classes, indicating a full decoupling between fluorescence efficiency and realized carbon gain.

Intermediate relationships, such as SIFyield-Gs, also weakened with rising O₃, suggesting that stomatal regulation of fluorescence efficiency is partly maintained but insufficient to buffer stress. These findings highlight that accumulated O₃ not only weakens individual linkages but also fragments the entire chain from absorbed light → stomata → fluorescence → carbon gain.

By comparing these canopy-scale patterns with chamber experiments, we see consistent evidence that O₃ drives a progressive decoupling of functional indicators. While chambers revealed leaf-level strain through impaired ΦPSII relationships, the remote sensing analysis demonstrates that the same disruption propagates across multiple efficiency metrics at the regional scale. This underscores the value of SIF and derived quantities like SIFyield for identifying when stress pushes canopies from coordinated functioning into a decoupled strain phase.

## 5. Conclusions

This study links leaf-level chamber experiments with regional satellite observations to advance mechanistic understanding of how O₃ affects soybean physiology under realistic exposure conditions. Across both scales, we find that the timing of O₃ stress is more important than its cumulative dose. In chambers, the first fumigation event caused the strongest suppression of photosynthesis (A and Gs) and efficiency (ΦPSII), initiating a strain phase from which plants did not fully recover. Recovery between weekly exposures was incomplete, leading to sustained growth suppression – despite the quite moderate levels of air pollution used herein.

The controlled chamber experiments applied repeated exposures to mildly elevated O₃, consistent with conditions typical of North American agroecosystems. We note that fumigation levels were below the thresholds set by the U.S. EPA for human health. The National Ambient Air Quality Standard is based on an 8-hour average of 70 ppb, while we show that exposures equivalent to a weekly 8-hour average of 40 ppb, or a 4-hour average of 80 ppb, were sufficient to cause significant stress and clear reductions in crop productivity. Our findings highlight the need for regulators to consider agricultural impacts when managing O₃ precursors, which is particularly urgent given that wildfire smoke, rising temperatures, and other climate-related drivers are likely to increase regional O₃ episodes in the future (Pusede et al., 2015; Xu et al., 2021).

At the regional scale, multi-year satellite observations confirmed that early-season O₃ episodes produced stronger declines in SIF, GPP, and Gs than equivalent late-season exposures, with recovery lagging for several weeks. Seasonal yield proxies were best explained not by total O₃ accumulation, but instead by exposures during early and peak growth phases, when canopies are expanding and reproductive sinks are being established. These findings bridge controlled and field conditions, demonstrating that even subtle, episodic O₃ events can constrain soybean productivity when they occur at vulnerable stages – highlighting the potential of functional indicators such as SIF to detect stress before irreversible damage occurs.

A limitation of this study is the coarse resolution of the TROPOMI SIF product (∼7 × 3.5 km), which restricts application to small fields. Nevertheless, the physiological mechanisms identified, and detection strategies developed herein, are broadly transferable across agricultural systems. The upcoming ESA FLEX mission, dedicated to fluorescence monitoring at finer spatial scales, carries potential to enhance the applicability of these approaches, enabling earlier and more precise detection of O₃ stress. Additionally, while stricter regulation of O₃ precursors in agricultural regions could help protect crop productivity, such measures are difficult to implement as high O₃ events often coincide with heat waves and drought, which themselves increase precursor emissions. We therefore emphasize the need to reduce ground-level O₃ as part of integrated strategies for crop health and global food security in a changing climate.

## Acknowledgements

This paper and related research have been conducted within the framework of the Italian National PhD Program in Earth Observation funded by the European Union – Next Generation EU through the Project of Italian Recovery and Resilience Plan (NRRP). The TROPOSIF products were generated by the TROPOSIF team conducted by NOVELTIS under the European Space Agency (ESA) Sentinel-5p+ Innovation activity Contract N° 4000127461/19/I-NS. We acknowledge funding from a National Science Foundation AGS Postdoctoral Research Fellowship (MR; 2306215). We would like to acknowledge the help of Lydia Tonnesen and Alyssa Belanger, who assisted with planting and caring for our soybean plants throughout the entire plant-level experiment.

