## Supplementary Information for "Mechanisms of Ozone Effects on Plant Stress in Soybean Across Growing Season: From Leaf to Regional Perspective"

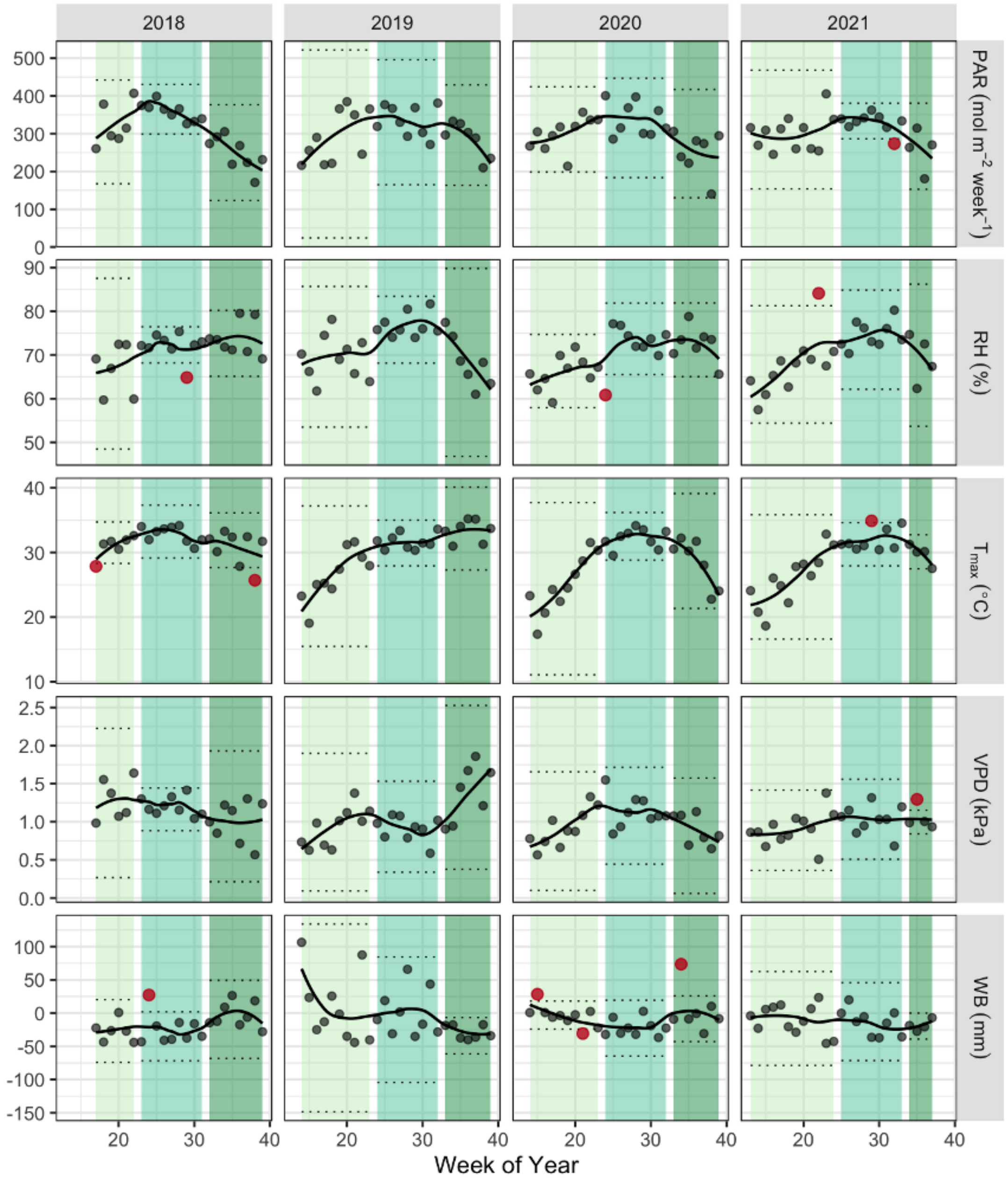


**Figure S1.** Weekly time series of environmental variables (PAR, RH, $T_{max}$ , VPD, and WB) in soybean fields in Crittenden County, Arkansas, across four growing seasons (April-September, 2018-2021). Points show weekly observations, and lines represent smoothed seasonal signatures. Shaded bands indicate the early (light green), peak (mid green), and late (dark green) phases of the growing season. Solid (non-transparent) points mark flagged outliers within each phase.

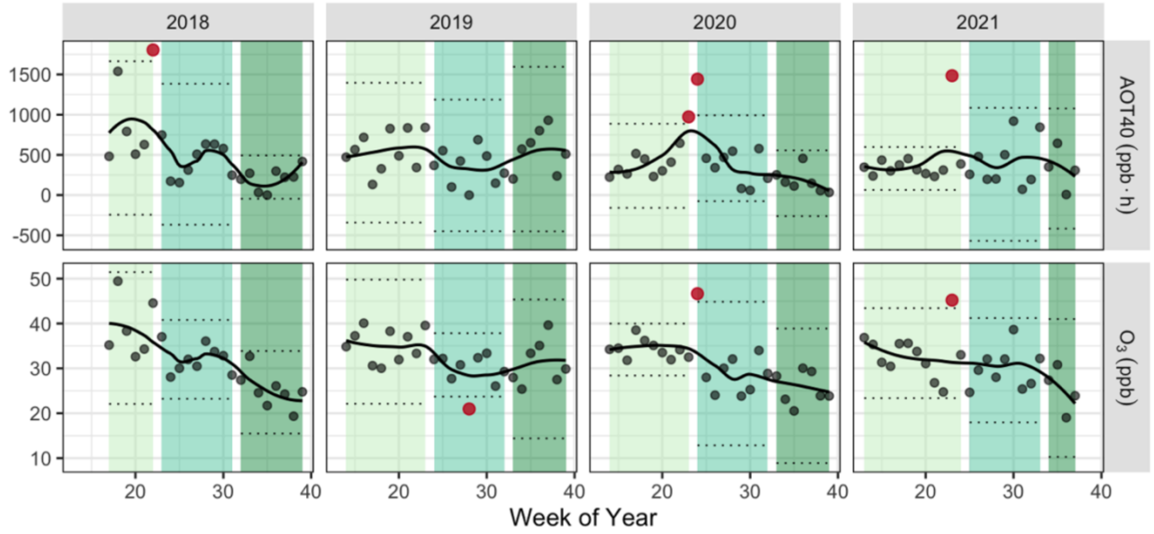


**Figure S2.** Weekly time series of AOT40 and mean $O_3$ in soybean fields in Crittenden County, Arkansas, across four growing seasons (April-September, 2018-2021). Points show weekly observations, and lines represent smoothed seasonal signatures. Shaded bands indicate the early (light green), peak (mid green), and late (dark green) phases of the growing season. Solid (non-transparent) points mark flagged outliers within each phase.

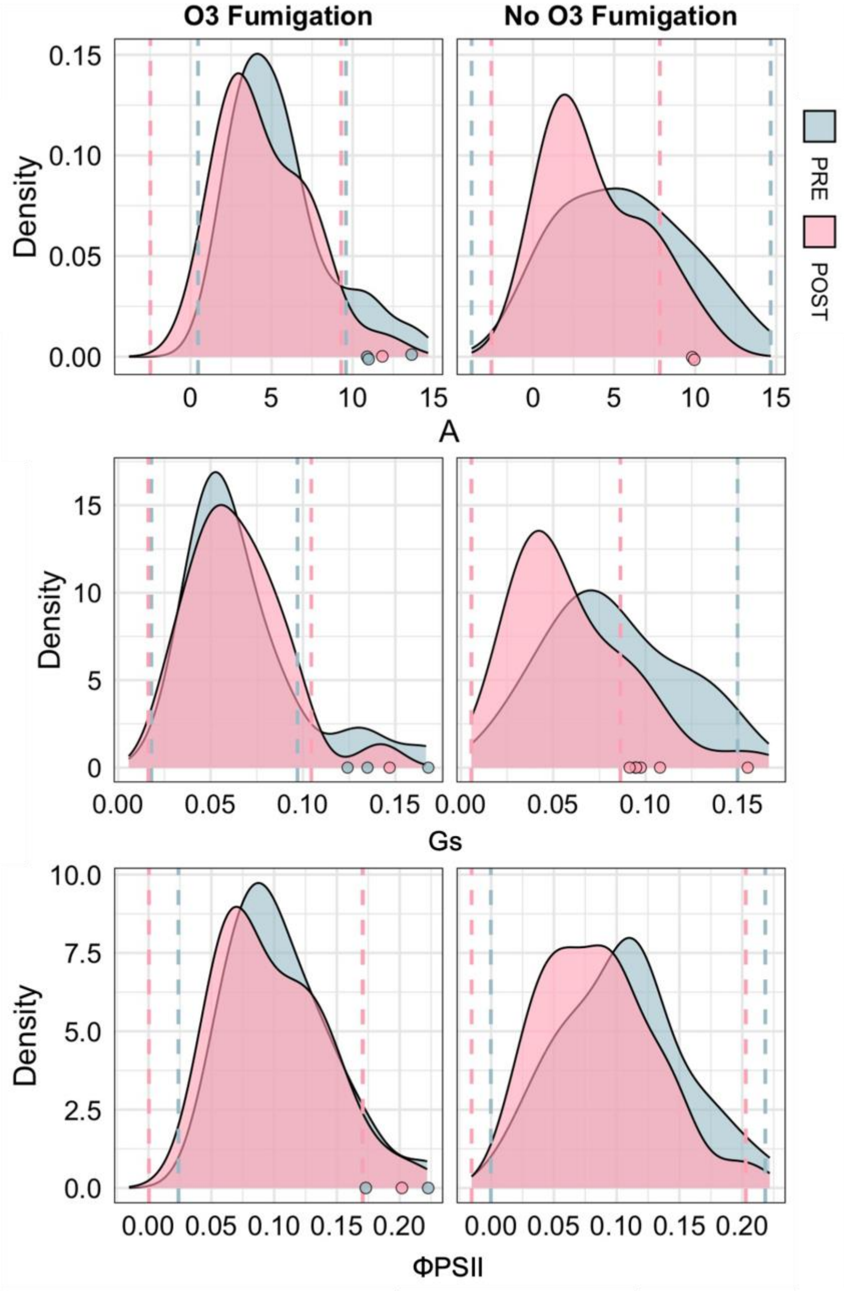


**Figure S3.** Distribution of pre- and post-fumigation leaf-level measurements of A (top), Gs (middle), and ΦPSII (bottom) for $O_3$-fumigated (left column) and control (right column) soybean plants across the five-week experiment. Dashed vertical lines indicate three-MAD thresholds used for outlier detection which are shown as circles.

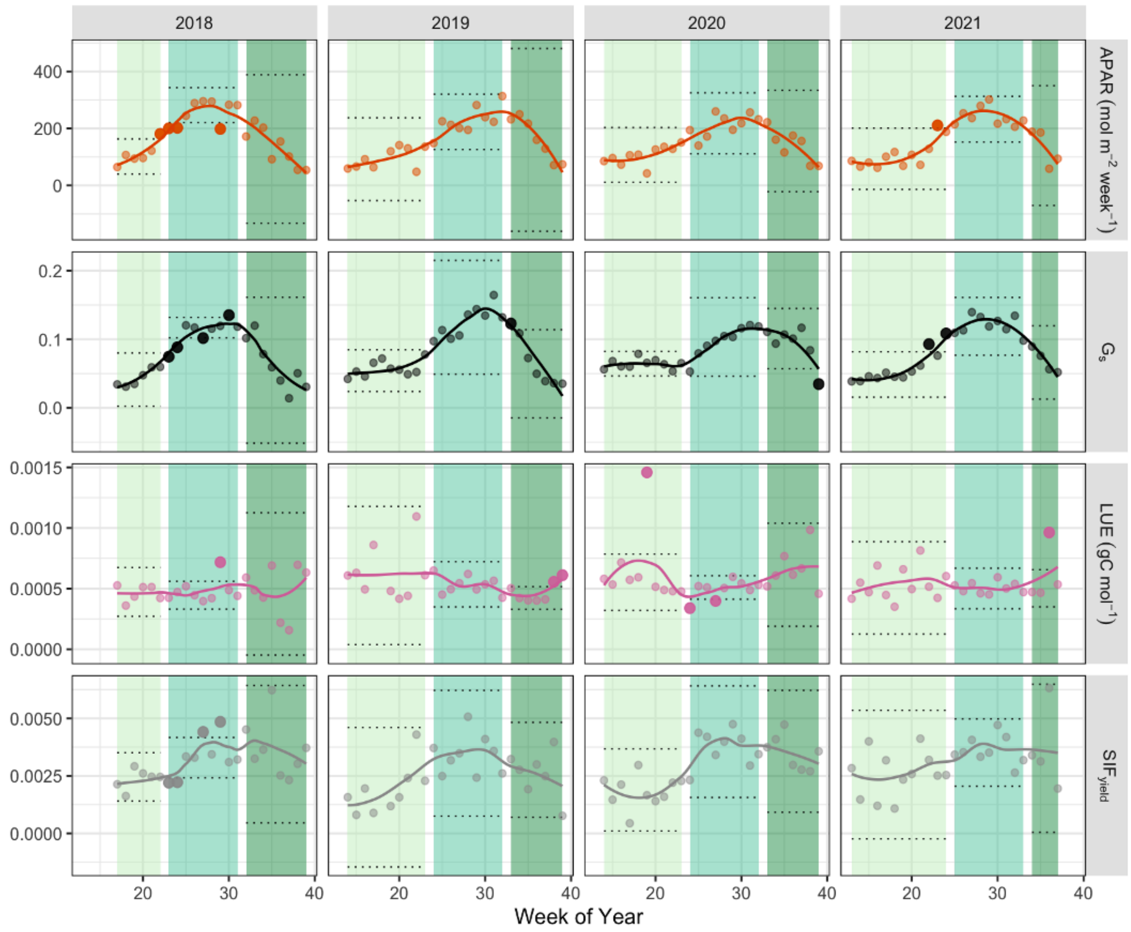


**Figure S4.** Weekly time series of additional vegetation indicators in soybean fields in Crittenden County, Arkansas, across four growing seasons (April-September, 2018-2021). Points show weekly observations, and lines represent smoothed seasonal signatures. Shaded bands indicate the early (light green), peak (mid green), and late (dark green) phases of the growing season. Solid (non-transparent) points mark flagged outliers within each phase.

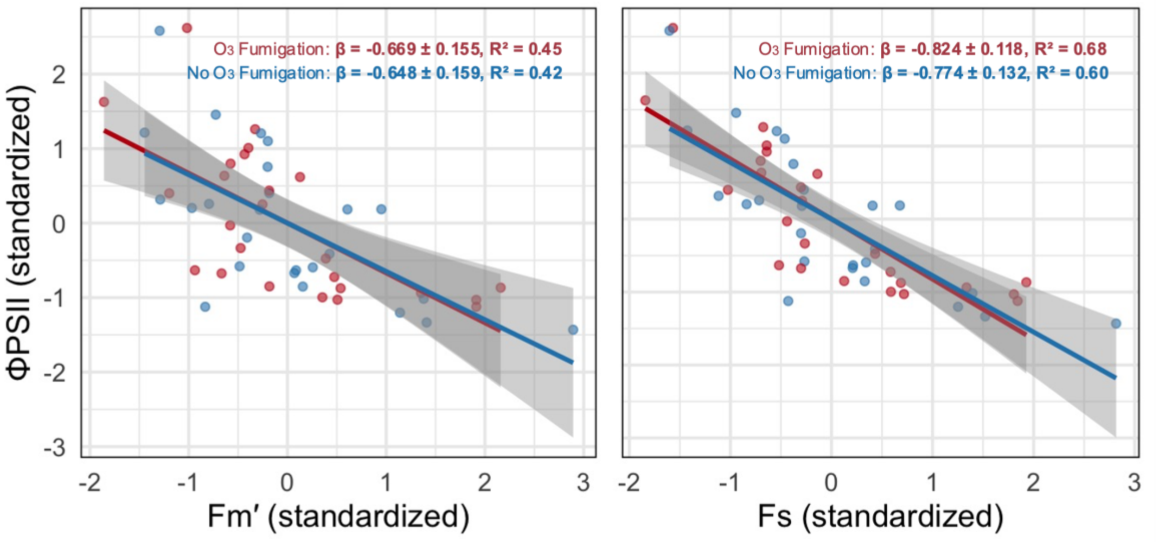


**Figure S5.** Relationships between ΦPSII and fluorescence components (Fm' - left, and Fs - right ) under $O_3$ fumigation (red) and control conditions (blue) presented by immediate post-chamber measurements across five weeks. Slopes (β ± SE) and $R^2$ are shown for each treatment; p-values < 0.001.

**Table S1.** Identified first and later $O_3$ episodes (ΔAOT40 > 350 ppb·h) in each of the four soybean growing seasons (2018-2021).

| Year | Timing | Week number |
|---|---|---|
| 2018 | First | 18 |
| 2018 | Later | 22 |
| 2019 | First | 19 |
| 2019 | Later | 23 |
| 2020 | First | 24 |
| 2020 | Later | 31 |
| 2021 | First | 23 |
| 2021 | Later | 30 |

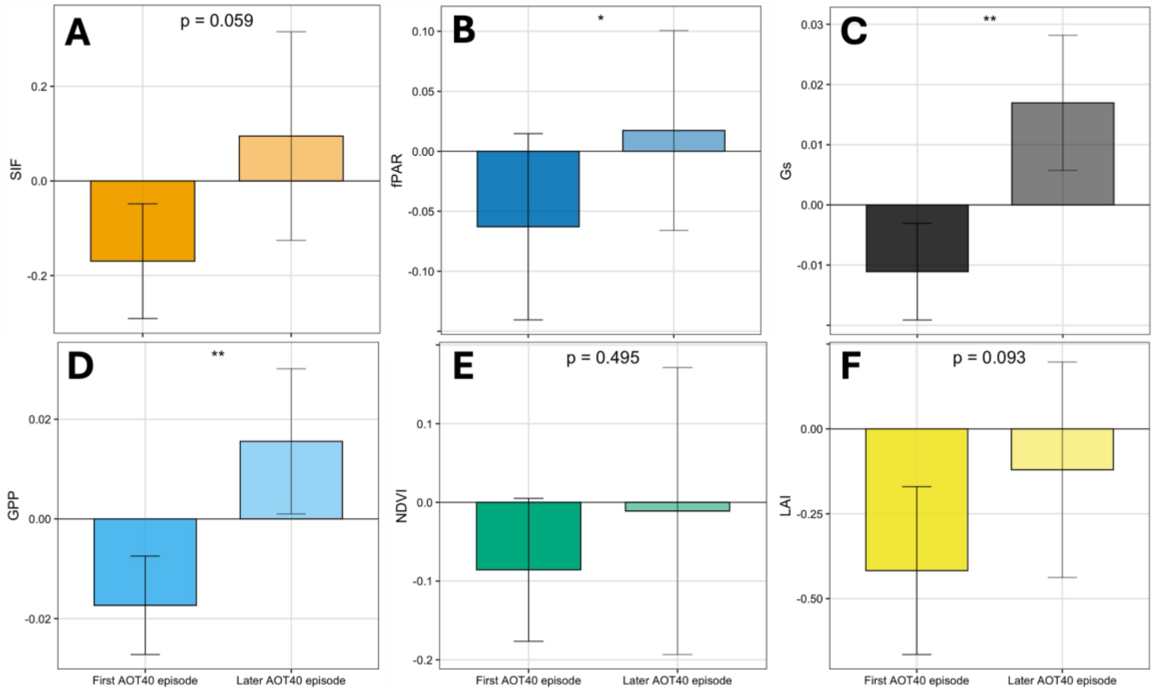


**Figure S6.** Residuals of selected vegetation indicators compared between the first and later $O_3$ episodes of equivalent AOT40 increase (>350 ppb·h) within each growing season (2018-2021). Bars represent mean ± SE across four paired episodes. Asterisks indicate significance levels (* $p < 0.05$).

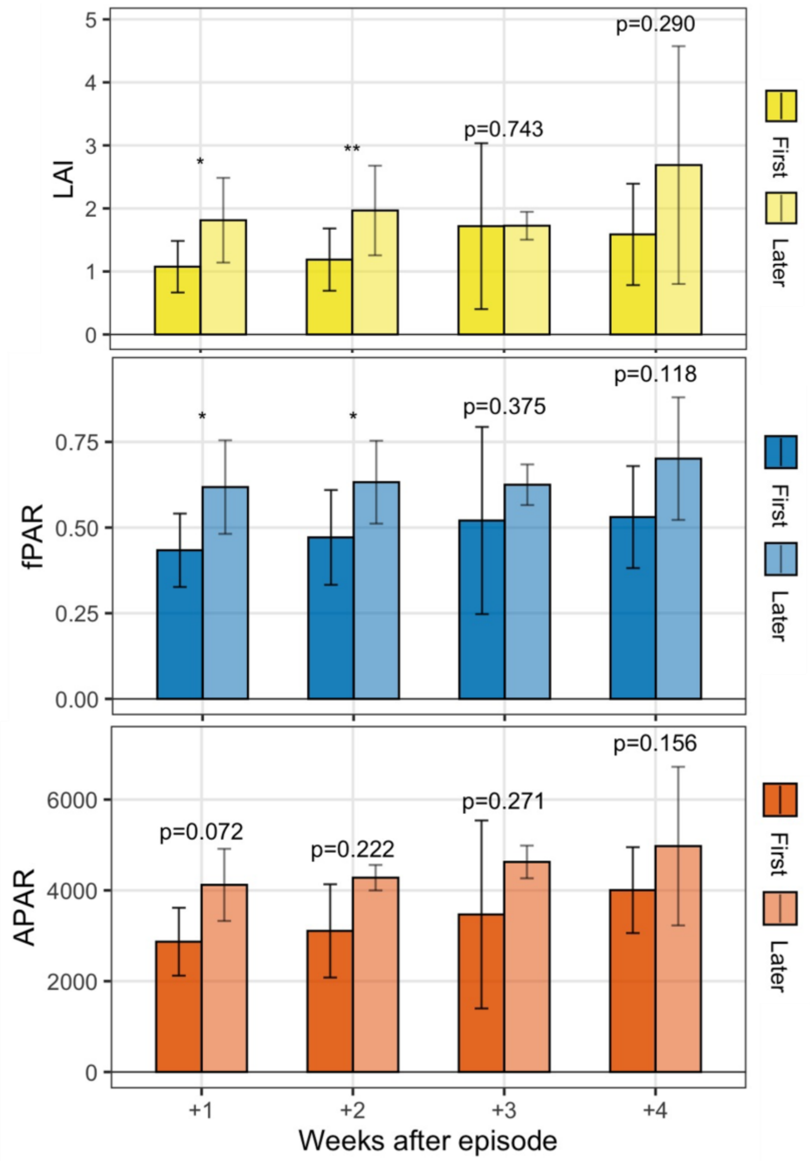


**Figure S7.** Recovery of structural vegetation indicators (LAI, fPAR, and APAR) following the first (dark bars) vs. later (light bars) seasonal $O_3$ episodes ($\Delta$AOT40 > 350 ppb·h). Bars show mean ± SE across four growing seasons (2018-2021). Asterisks indicate significance levels (* $p < 0.05$).

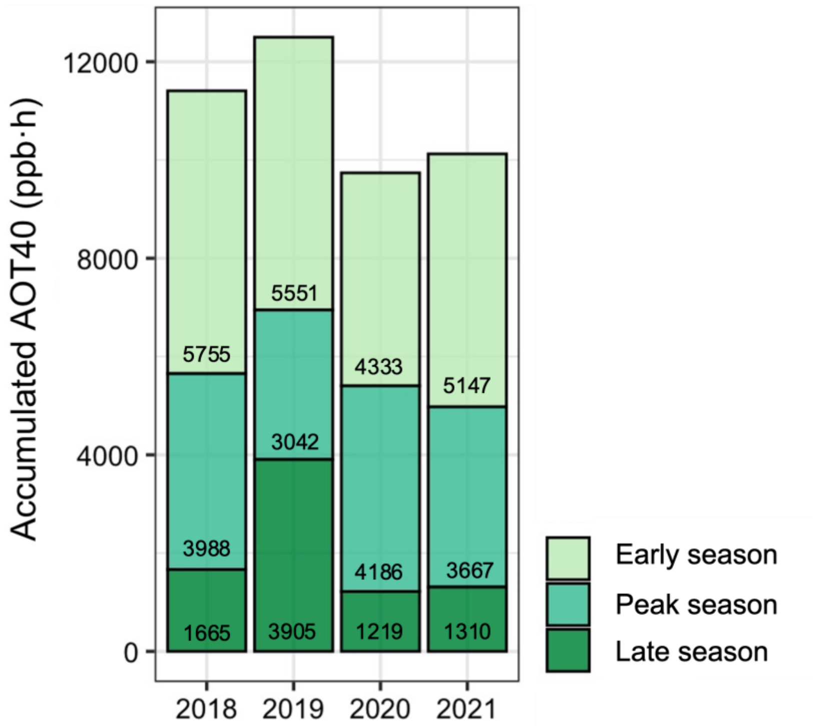


**Figure S8.** Accumulated AOT40 (ppb·h) in soybean fields in Crittenden County, Arkansas, across four growing seasons (April-September, 2018-2021) divided by growing season phases (early, peak, late).

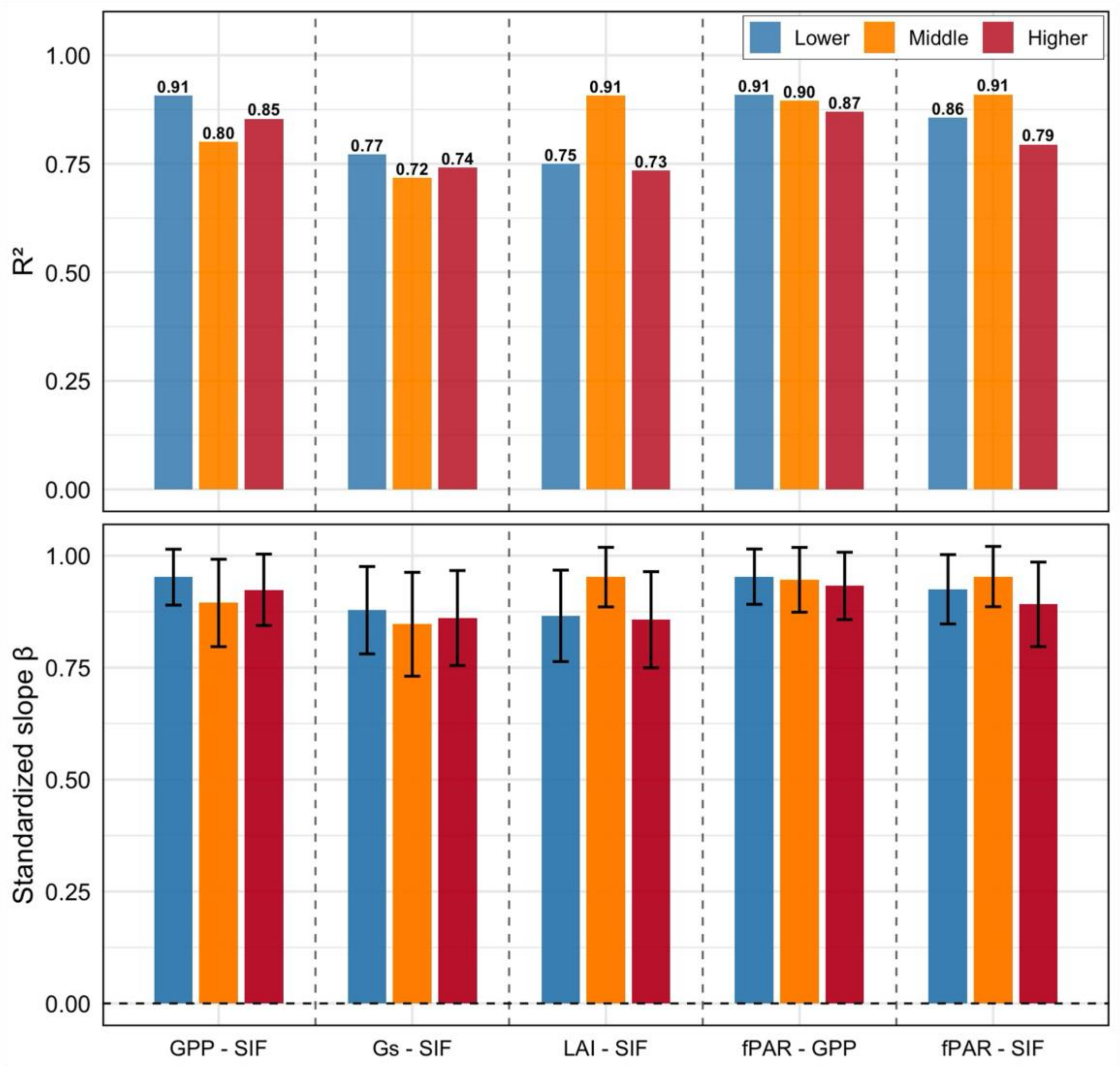


**Figure S9.** Relationships between vegetation health indicators during the early and peak soybean growing season by different $O_3$ exposure. Bars show standardized regression slopes (β, bottom) ± SE, and coefficients of determination ($R^2$, top) for pairs of functional and structural indicators. Lower (blue), middle (orange), and higher (red) tertiles correspond to AOT40 thresholds of ≤311, 311-508, and ≥508 ppb·h, respectively.
